# TBC1D15 potentiates lysosomal regeneration from damaged membranes

**DOI:** 10.1101/2022.12.14.520480

**Authors:** Anshu Bhattacharya, Rukmini Mukherjee, Santosh Kumar Kuncha, Melinda Brunstein, Rajeshwari Rathore, Stephan Junek, Christian Münch, Ivan Dikic

## Abstract

Acute lysosomal membrane damage reduces the cellular population of functional lysosomes. However, these damaged lysosomes have a remarkable recovery potential independent of lysosomal biogenesis and remain unaffected in TFEB/TFE3-depleted cells. We combined proximity labelling based proteomics, biochemistry and high-resolution microscopy to unravel a new lysosomal membrane regeneration pathway which is dependent on ATG8, lysosomal membrane protein LIMP2, the Rab7 GAP TBC1D15, and proteins required for autophagic lysosomal reformation (ALR) including Dynamin2, Kinesin5B and Clathrin. Upon lysosomal damage, LIMP2 act as a lysophagy receptor to bind ATG8, which in turn recruits TBC1D15 to damaged membranes. TBC1D15 hydrolyses Rab7-GTP to segregate the damaged lysosomal mass and provides a scaffold to assemble and stabilize the ALR machinery, potentiating the formation of lysosomal tubules and subsequent Dynamin2-dependent scission. TBC1D15-mediated lysosome regeneration was also observed in a cell culture model of oxalate nephropathy.

## Main

Lysosomes are single-membrane-bound organelles found in eukaryotic cells. They are the major site for degradation of macromolecules in the endolysosomal system and represent the terminal point of the autophagic pathway. This function is well characterized, but recent studies have shown that lysosomes are also involved in intracellular signalling, metabolite sensing, and antigen presentation (Saftig and Puertollano, 2021; Bonam et al., 2019). During bulk and selective autophagy, autophagosomes containing damaged organelles, proteins and pathogens fuse with lysosomes to form autolysosomes. The low pH of autolysosomes activates luminal cathepsins which can then break down the autophagic cargo.

The acute loss of lysosomal function can result from lysosomal membrane damage, triggered by mineral crystals such as silica and monosodium urate, toxins, lipids,β-amyloid, and lysosomotropic compounds. The resulting inflammatory processes contribute to neurodegenerative disorders such as Parkinson’sdisease, and induce kidney dysfunction as seen in hyperuricaemic nephropathy (Emmerson et al., 1990; Boya and Kroemer, 2008; Dehay et al., 2010). Lysosomal membrane damage triggers various cellular responses depending on the severity. Small perturbations in lysosomal membranes cause the recruitment of endosomal sorting complex required for transport (ESCRT) machinery, which repairs membranes in a calcium-dependent manner (Skowyra et al., 2018; Radulovic et al., 2018). Extensive damage causing lysosomal perforation promotes lysophagy, the selective autophagy of damaged lysosomes (Maejima et al., 2013; Papadopoulos and Meyer, 2017). Membrane damage also inactivates mammalian target of rapamycin (mTOR), which normally maintains transcription factor EB (TFEB) in an inactive state by phosphorylation. The dephosphorylation of TFEB causes nuclear translocation and the induction of genes controlling lysosomal biogenesis (Napolitano and Ballabio, 2016).

Membrane damage exposes the glycosylated luminal domains of lysosomal membrane proteins, resulting in the recruitment of galectins (mainly galectins 1, 3, 8 and 9). Galectins facilitate the assembly of protein complexes that coordinate repair and autophagy at the site of damage (Jia et al., 2020). Several ubiquitin E3 ligases, including TRIM16 and the SCF ligase component FBXO27, are recruited to such sites (Yoshida et al., 2017). Damaged membranes are also marked by specific ubiquitin chain signatures, including K63 ubiquitin chains (Koerver et al., 2019). The AAA-ATPase p97 (VCP) and the deubiquitinating enzyme YOD1 target ubiquitinated membrane proteins for proteasomal degradation (Papadopoulos et al., 2017). Ubiquitinated membrane proteins bind to autophagy receptors such as NDP52, Optineurin, CALCOCO2, p62 and TAX1BP1. Quantitative proteomics and proximity biotinylation analysis of autophagy adaptors, cargo receptors, and galectins responding to acute lysosomal damage have identified TAX1BP1 and the kinase TANK1-binding kinase 1 (TBK1) as important regulators of lysophagic flux (Eapen et al., 2021). Ubiquitination also mediates the recruitment of FIP200, a component of the ULK1 autophagosome initiation complex, which initiates the formation of a phagophore (Fujita et al., 2013). TRIM16 binds ATG16L to facilitate the maturation of the autophagosome at the site of damage (Chauhan et al., 2016).

Prolonged activation of starvation induced autophagy activates autophagic lysosome reformation (ALR), in which membranes derived from autolysosomes are recycled into new lysosomes. ALR is induced by prolonged autophagy. During this process autolysosomes extrude tubular membrane structures that undergo scission to generate proto-lysosomes, which subsequently mature into functional lysosomes (Yu et al., 2010). The tubular structures are formed when phosphatidylinositol 4-phosphate [PI(4)P] is converted to phosphatidylinositol 4,5-bisphosphate [PI(4,5)P2] by the sequential actions of PI-4 and PI(4)P-5 kinases. Microdomains enriched in PI(4,5)P2 then recruit effector proteins such as CLTC, which promote budding and the initiation of tubules. The tubules grow, which requires the microtubule-associated kinesin motor protein KIF5B, before scission induced by Dynamin2/DNM2 (Chen and Yu, 2017).

In this study, we identified a novel mechanism in which functional lysosomes are regenerated from damaged lysosomal membranes. We found that lysosomes damage induced by the lysosomotropic compound L-leucyl-L-leucine methyl ester (LLOMe),(Thiele and Lipsky, 1990), could recover ∼50% of their original activity independently of lysosome biogenesis. This process is coordinated by the Rab7 GAP (GTPase activating protein) TBC1D15, which binds to LC3 on damaged lysosomes. Tre2–Bub2–Cdc16 (TBC) domain proteins such as TBC1D15 are negative regulators of the Rab family of small GTPases (Onoue et al., 2013; Wong et al., 2018). They contain a highly conserved TBC domain, one or more LC3 interaction region (LIR) motifs, and can interact with ATG8 proteins (Popovic et al., 2012). We found that the lysosomal membrane protein LIMP2 binds to ATG8 proteins upon LLOMe treatment. This stabilizes TBC1D15 on damaged lysosomes and makes the organelles deficient for Rab7, segregating them from vesicular traffic. Proximity labelling of TBC1D15 and LAMP1 identified DNM2, KIF5B and CLTC as important players in the regeneration of lysosomes from damaged membranes. Mutating the N-terminal domain of TBC1D15 abolished the interaction with DNM2, indicating that TBC1D15 is a scaffold for the assembly of the lysosomal regeneration machinery.

## Results

### Lysosomal recovery following LLOMe treatment involves sequential TFEB-independent and TFEB-dependent phases

We treated HeLa cells with control TFEB or TFE3 siRNA for 48 h, then with 1 mM LLOMe for 2 h to disrupt lysosomal membranes. Knockdown of TFEB and TFE3 prevents lysosomal biogenesis. This was followed by a washout in LLOMe-free medium. We then added 100 nM of the pH-sensitive LysoTrackerRed probe for 30 min before fixation and used confocal microscopy to detect functional (acidic) lysosomes. LLOMe treatment reduced the LysoTracker signal intensity by ∼80%, but the cells recovered to ∼50% of the original signal intensity within 2 h of LLOMe washout. Recovery during the first 2 h was similar in control cells and in cells treated with TFEB or TFE3 siRNA, but control cells accumulated a larger population of acidic lysosomes than TFEB/TFE3-depleted cells after a longer recovery period of 6–8 h (**Fig 1A, S1A**).

**Figure 1:**
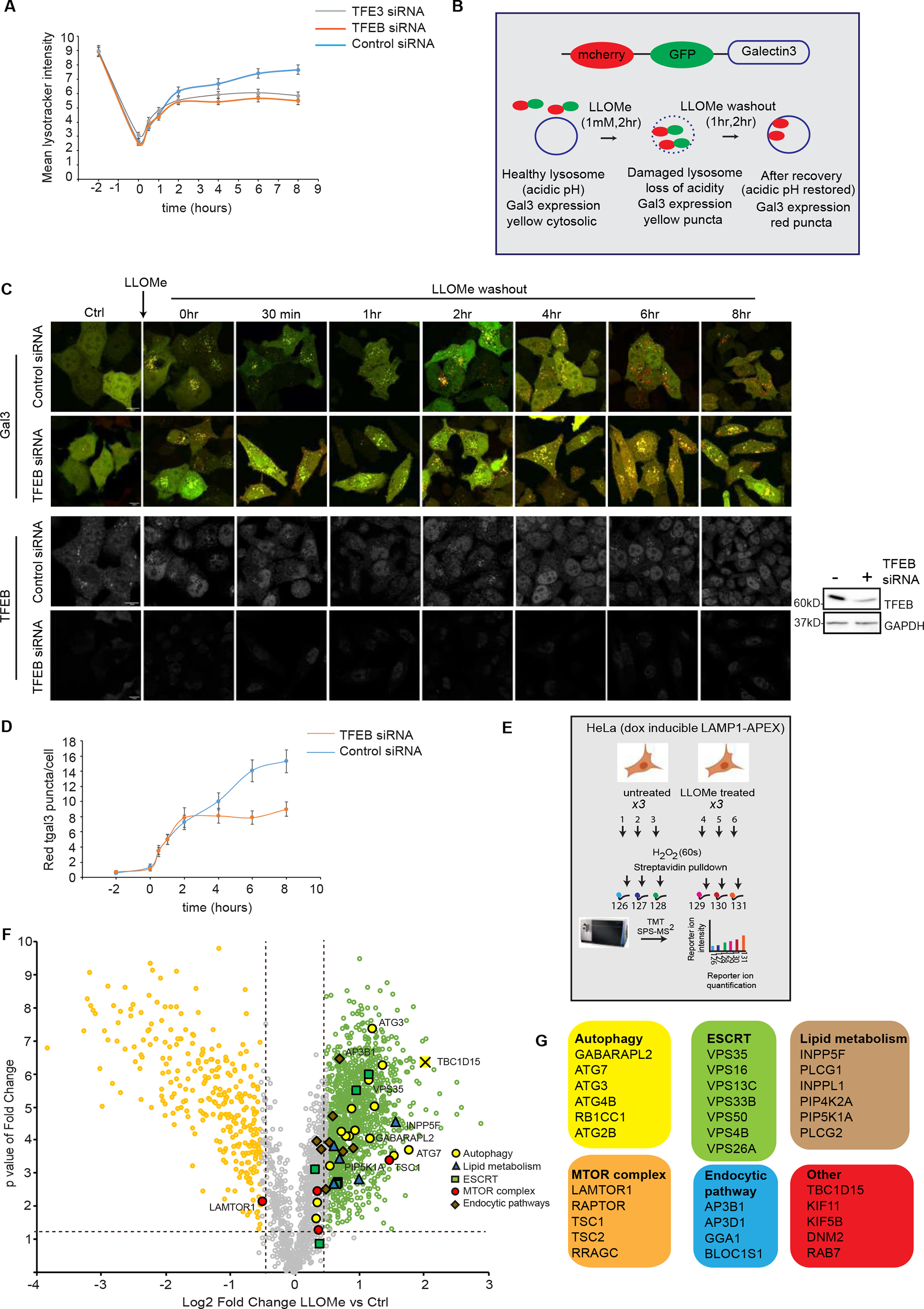
Cellular response to LLOMe-mediated acute lysosomal damage. a) HeLa cells were treated with control, TFEB or TFE3 siRNA for 48 h. Lysosomal damage was induced by 1mM LLOMe for 2 h. The LLOMe containing media was then replaced with fresh media to allow recovery. Cells were fixed at different time points after LLOMe washout. LysoTracker Red was loaded 30 min before the cells were fixed for confocal imaging. LysoTracker Red intensity was measured at different time points before and after LLOMe washout. Data are means ± SEM of 60 cells taken from three independent experiments. b) Schematic of the RFP GFP dual-tagged Gal3 (tfGal3) reporter construct used for the lysosomal regeneration flux assay. c) HeLa cells with stable expression of tfGal3 were treated with control or TFEB siRNA for 48 h. Lysosomal damage was induced by treating cells with LLOMe (1mM, 2 h). Cells were cultured in LLOMe-free medium for different durations similar to panel (a). The number of red Galectin3 puncta was counted at each time point before and after LLOMe washout. TFEB knockdown efficiency is noted by western blotting. d) Graph showing numbers of red Galectin3 puncta per cell obtained from the experiments in panel (c). Data are means ± SEM of 50 cells taken from three independent experiments. e) Schematic of the LAMP1-APEX proximity labelling assay. f) Volcano plot to compare the biotinylated landscape of APEX2-tagged LAMP1 with and without treatment with 1mM LLOMe for 2 h. Green and yellow circles indicate biotinylated proteins that are significantly enriched and significantly depleted in LLOMe-treated cells, respectively. g) Major categories and candidate proteins enriched by LLOMe treatment that are relevant to this study following pathway analysis in Perseus.

The extent of lysosomal damage and recovery was also assessed using a HeLa reporter cell line with stable expression of RFP–GFP (tandem fluorescent)-tagged Gal3 (tfGal3). This probe is based on the principle that GFP fluorescence is attenuated by the acidic lysosomal environment whereas RFP fluorescence is comparatively stable, so that damaged lysosomes are marked by yellow puncta whereas recovered, acidic lysosomes appear red (**Fig 1B**). Control and TFEB-depleted HeLa cells expressing tfGal3 were treated with 1 mM LLOMe for 2 h followed by washout as above. LLOMe treatment led to the formation of yellow tfGal3 puncta, indicating lysosome damage, but washout increased the number of red Gal3 puncta, indicating recovery. As discussed above for LysoTracker Red intensity, the formation of red Gal3 puncta was independent of TFEB/TFE3 during the first 2 h, but control cells again showed a greater degree of recovery than TFEB-depleted cells at later time points of 6–8 h (**Fig 1C, 1D**).

To understand whether the newly formed lysosomes have active hydrolases, we used the fluorogenic substrate of lysosomal hydrolases DQ-BSA. DQ-BSA enters the cell through the endolysosomal pathway and reaches the lysosomes, where it is cleaved by lysosomal proteases, resulting in the de-quenching of the dye which is visualized as a bright fluorescence in cells. In control siRNA treated cells, treatment with LLOMe led to a sharp decline in DQ-BSA fluorescence to < 20% of the intensity measured in untreated cells. Washout of LLOMe for 2 h restored the hydrolase activity to ∼60% of the activity measured in untreated cells. TFEB depleted cells had lower intensities of DQ-BSA under all conditions indicating lysosomal activity was compromised in these cells even without LLOMe treatment. However, LLOMe washout led to increase of DQ-BSA fluorescence following a pattern which was similar to that observed in cells treated with control siRNA (**Fig. S1B, S1C**). This indicated that, while lysosomal hydrolase synthesis was dependent on active TFEB, the observed fluorescence recovery in the first 2 hours of LLOMe washout was not directly influenced by TFEB. The initial TFEB independent recovery of hydrolase activity may be explained by previous reports which demonstrate that 1mM LLOMe treatment for 2 h destabilises the lysosomal membrane causing lysosomal membrane permeabilisation (LMP) but does not release large molecules of size >10 kDa (like cathepsins) from lysosomes (Repnik et al, 2017).

These experiments indicated that LLOMe-mediated lysosomal damage is followed by a biphasic recovery profile, comprising a rapid initial phase (1–2 h after LLOMe washout) that is independent of TFEB/TFE3 and a subsequent phase (6–8 h after LLOMe washout) that is dependent on regulators of lysosomal biogenesis.

### Lysosomal recovery following LLOMe treatment is independent of mTOR

Since mTOR (mammalian target of Rapamycin) is an important determinant of lysosomal biogenesis, we tested the activity of mTOR in LLOMe treated cells and during the lysosomal recovery period following LLOMe removal. Control cells have high levels of phospho-p70 S6K (Thr 389), indicating active mTOR signalling. LLOMe treatment led to almost complete loss of mTORC1 activity. Lysosomal recovery in LLOMe free media for at least 4 h was needed to reactivate mTORC1 mediated phosphorylation of p70 S6 kinase (Fig. S1D). The initial phase of lysosomal recovery appears to be independent of mTORC1 activation. The nuclear localization of TFEB in cells in Fig S1A in the first period of LLOMe recovery also indicates reduced mTORC1 activity in the first 2 h of LLOMe recovery.

### The biotin-labelled LAMP1 interactome is modified significantly by membrane damage

We investigated the rapid phase of lysosomal recovery in more detail using a proximity labelling assay. HeLa cells expressing doxycycline-inducible LAMP1-APEX2 were treated with LLOMe for 2 h, followed by exposure to biotin and H_2_O_2_ and the analysis of biotin-labelled proteins by mass spectrometry (MS). HeLa cell extracts were analysed by 6-plex tandem mass tag (TMT) labelling, revealing significant differences in the biotin-labelled LAMP1 interactome of the LLOMe-treated and control cells (**Fig 1E**). We identified several classes of differentially represented proteins, including those involved in autophagy (RB1CC1, GABARAPL1, ATG3 and ATG7), the mTOR pathway (LAMTOR1, TSC1, TSC2 and RAPTOR), the ESCRT pathway (VPS35, VPS16, VPS13C and VPS50) and lipid metabolism (PIP4K2A, PIP5K1A, INPPL1, INPP5F and PLCG2) as well as other proteins such as BLOC1S1, TBC1D15, Rab7, DNM2 and kinesins (**Fig 1F, 1G**).

### LC3-binding proteins are recruited to lysosomes following membrane damage

ATG8 proteins are recruited to damaged lysosomes, so we also characterised the interactome of GST-tagged LC3B following LLOMe treatment. HEK 293T cells were lysed following treatment with 1 mM LLOMe (untreated controls were lysed in parallel) and the lysates were incubated with purified GST-LC3B in a pulldown assay. GST-LC3B interaction partners were analysed by label-free quantification (LFQ)-based MS. We found that TBC1D15 was significantly enriched in the GST-LC3 pulldown from the LLOMe-treated cells (**Fig 2A**). This protein was also enriched in the biotin-labelled interactome of LAMP1-APEX2 but only in the LLOMe-treated cells (**Fig 1F**). Western blotting of biotin-labelled proteins following streptavidin pulldown suggested that TBC1D15 is recruited to LAMP1^+^ damaged lysosomal membranes after LLOMe treatment, but LLOMe washout for 2 h reversed this recruitment. LC3 was also recruited to LAMP1 following LLOMe treatment and remained associated with LAMP1 2 h after LLOMe washout (**Fig 2B**). Therefore, the TBC1D15–LAMP1 interaction was dynamic, and may have an important role in the first phase of lysosomal recovery following membrane damage. TBC1D15 was also identified as part of the biotin-labelled LC3B interactome in LLOMe-treated cells in a recent study of lysophagy (Eapen et al., 2021). To confirm our proteomics data, we treated HEK 293T cells expressing hemagglutinin (HA)-tagged TBC1D15 with 1 mM LLOMe, lysed the cells at different time points after treatment, and incubated the lysates with anti-HA beads to recover HA-TBC1D15 by immunoprecipitation. A time course analysis revealed that the TBC1D15–LC3B interaction increased during the LLOMe treatment (**Fig 2C**). Taken together, these findings suggest that TBC1D15 is recruited to damaged lysosomes following LLOMe-mediated membrane damage, and interacts with ATG8 proteins.

**Figure 2:**
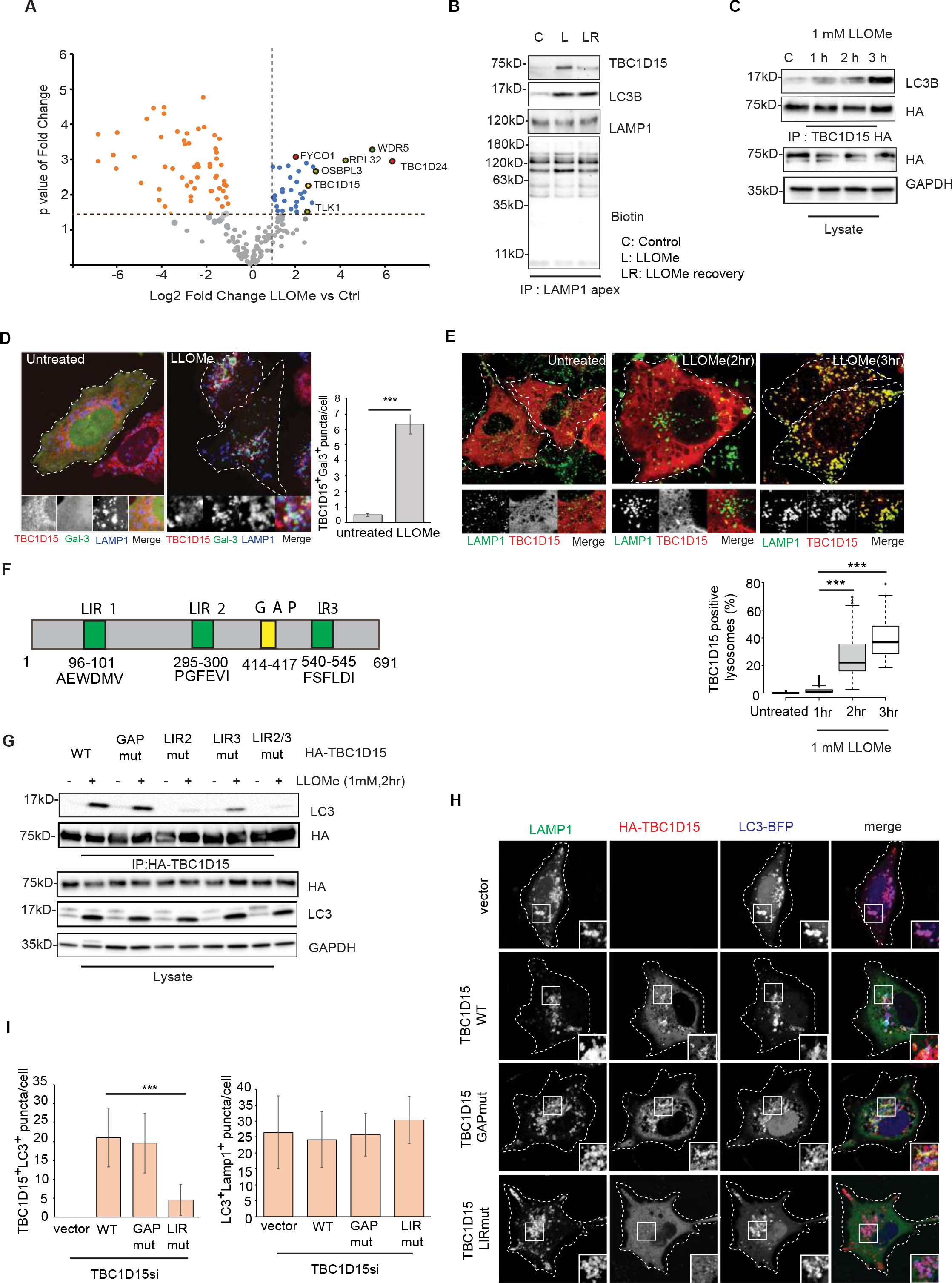
TBC1D15 is recruited to damaged lysosomes in an ATG8-dependent manner. a) GST-LC3B was incubated with cell lysates from control or LLOMe-treated cells followed by the detection of interacting proteins by MS. The volcano plot shows LC3-binding proteins enriched after treating cells with LLOMe. b) Cells expressing doxycycline-inducible LAMP1-APEX were treated with LLOMe to damage lysosomes and LLOMe recovery was performed by changing the medium. 500 µM biotin-tyramide was added during the last hour of LLOMe treatment or recovery period and then cells were exposed to 1 mM H_2_O_2_ for 1 min for inducing biotinylation. Biotinylated proteins were pulled down using streptavidin-resin. Western blots of samples representing different conditions were probed with antibodies to detect the proteins of interest (TBC1D15, LC3B and LAMP1). C = control. L = treatment with 1 mM LLOMe for 2 h. LR = recovery of 2 h after LLOMe treatment. c) HEK 293T cells expressing HA-TBC1D15 were treated with 1 mM LLOMe for 1, 2 or 3 h. LC3B bound to TBC1D15 under each condition was assessed by immunoprecipitation using HA beads followed by western blots. d) HeLa cells were treated with1mM LLOMe for 2 h followed by staining for endogenous TBC1D15 (red), LAMP1 (blue) and GAL3 (green) using specific antibodies. Data are means ± SEM of 50 cells taken from three independent experiments and were counted for determining the Gal3^+^ TBC1D15^+^ puncta/cell(***p≤0.001). e) HeLa cells were transfected with HA-TBC1D15 and treated with 1 mM LLOMe for different 2 h or 3 h. Endogenous LAMP1 was stained along with HA-TBC1D15 to assess the degree of TBC1D15 recruitment to lysosomes. Data are means ± SEM of at least 100 cells taken from three different experiments for each time point (***p≤0.001). f) Domain architecture of TBC1D15 showing potential LIR motifs (predicted by screening the iLIR database) and the known GAP motif. g) HEK 293T cells expressing wild-type or HA-tagged mutants of TBC1D15 were treated with 1mM LLOMe for 2 h or left untreated and then tested by HA pulldown to check the effect on LC3B binding. h) HeLa cells were treated with TBC1D15 siRNA for 48 h followed by transfection with constructs encoding HA-tagged wild-type or mutant TBC1D15 and LC3-BFP. Cells were treated with 1 mM LLOMe for 2 h, fixed, stained for endogenous LAMP1 and HA, and imaged by confocal microscopy to assess the extent of TBC1D15 recruitment to damaged lysosomes. i) Images from the experiment in panel (h) were analysed using FIJI to evaluate the recruitment of TBC1D15 to LAMP1-labelled lysosomes and to check for the colocalization of LC3 and LAMP1. Data are means ± SEM of 40 cells from three independent experiments (***p≤0.001).

### TBC1D15 is recruited to damaged lysosomes following acute lysosomal damage

Next, we visualized the cellular distribution of endogenous TBC1D15 in LLOMe-treated HeLa cells by immunofluorescence analysis. The localization of TBC1D15 was primarily cytosolic in control cells, sometimes resembling structures which may be mitochondria but after LLOMe treatment for 2 h it was recruited to damaged (Galectin3-GFP positive) lysosomes (**Fig 2D**). Unlike ESCRTs, or Galectin3 which are recruited within 30 min-1 h of LLOMe-mediated lysosomal damage (and are involved in lysosomal repair), HA-TBC1D15 is recruited at a later stage (2-3 h after LLOMe treatment) (**Fig 2E**). HA-TBC1D15 also colocalized with LC3-BFP and Galectin3-GFP in LLOMe-treated cells (**Fig S2A**). TBC1D15^+^ lysosomes were also positive for ubiquitin in LLOMe-treated cells (**Fig S2B**). Treatment with the mTORC1 inhibitor Torin1, the CASM (conjugation of ATG8 to single membrane) inducer Monensin, serum starvation with Earle’s balanced salt solution (EBSS) or the treatment of cells with the mitochondrial proton gradient uncoupler carbonyl cyanide *m*-chlorophenylhydrazone (CCCP) did not induce the formation of TBC1D15 puncta (**Fig S2C**). There results suggest that TBC1D15 localizes specifically to damaged lysosomes, confirming the suitability of our proximity labelling strategy and interactome experiments.

### TBC1D15 recruitment to lysosomes involves two of three available LIR motifs

TBC proteins feature so-called LC3 interaction region (LIR) motifs (Popovic et al., 2012). Screening the TBC1D15 amino acid sequence against the iLIR database (Jacomin et al., 2016) revealed the presence of three such putative motifs (**Fig 2F**). To determine which of the motifs is involved in the response to LLOMe-mediated damage, we introduced multiple point mutations within each LIR motif of the HA-TBC1D15 protein, and overexpressed each mutant in HEK 293T cells. We then assessed the ability of the mutated TBC1D15 to bind LC3B by the immunoprecipitation of HA-TBC1D15 from the lysates of LLOMe-treated cells (**Fig 2G**). We found that mutating the LIR motif spanning amino acid residues 295–300 (TBC1D15[F297AV299AI300A], LIR2mut) led to a significant loss of LC3B binding. This particular motif has previously been identified as the most active LIR motif in TBC1D15 (Yamano et al., 2014). Mutating the LIR motif spanning amino acid residues 540–545 (TBC1D15[F540AF542AL543AI545A], LIR3mut) also caused some reduction in LC3B binding, and the double mutant (LIR2/3mut) lost the ability to interact with LC3B almost completely. In contrast, mutating the LIR motif spanning amino acids 96–101 (TBC1D15[W98AV101A]) had no effect on LC3B binding (**Fig S2D**). Given that TBC1D15 is a Rab7-GAP (Peralta et al., 2010), we also tested a GAP mutant (TBC1D15[D414AK417A], GAPmut) but observed no effect on LC3B binding in LLOMe-treated cells (**Fig 2G**). The interaction between TBC1D15 and GABARAPL2 was dependent on the same LIR motifs (**Fig S2E**).

Next, we determined the localization of the LIR and GAP mutants of TBC1D15 in LLOMe-treated cells. Endogenous TBC1D15 was depleted by exposing cells to the corresponding siRNA and then reconstituted with the HA-tagged wild-type or mutant versions of TBC1D15 (GAPmut and LIR2/3mut). Only the LIR mutant failed to localize with LLOMe-damaged lysosomes, whereas wild-type TBC1D15 and the GAP mutant showed robust recruitment (**Fig 2H, I**). Interestingly, the depletion of TBC1D15 did not reduce the percentage of LAMP1^+^ and LC3^+^ puncta after damage (**Fig 2I**), indicating that TBC1D15 does not affect LC3 recruitment to damaged LAMP1 membranes. These findings ruled out the possibility that TBC1D15 acts as a classical lysophagy receptor, in which case it would recruit ATG8 proteins to sites of damage. Instead, the presence of ATG8 proteins on damaged lysosomal membranes is necessary to recruit of TBC1D15 to damaged membranes.

### The GAP activity of TBC1D15 is stimulated by membrane damage

Given that TBC1D15 has both LIR and GAP functions, we determined the status of its GAP activity following lysosomal damage using a Rab-interacting lysosomal protein (RILP) pulldown assay to assess the amount of GTP-bound Rab7 present in cells treated with LLOMe compared to untreated controls. RILP specifically interacts with the GTP-bound form of Rab7 and is thus used as a bait to pull down Rab7-GTP from lysates (Sun et al., 2009). TBC1D15 GAP activity converts Rab7-GTP to Rab7-GDP, so the amount of Rab7-GTP precipitated with RILP can be used as a readout of TBC1D15 GAP activity.

In TBC1D15-depleted cells, the amount of Rab7-GTP was higher after lysosomal damage than in untreated cells. When reconstituted with wild-type TBC1D15, there was no difference in the amount of Rab7-GTP between LLOMe-treated cells and controls. Cells expressing the GAP mutant accumulated more of the GTP-bound Rab7 as expected. Interestingly, we also pulled down more Rab7-GTP from cells expressing the LIR mutant than from those expressing wild-type TBC1D15, indicating that GAP activity was also lower in the LIR mutant (**Fig S2F**). This suggested that LLOMe-mediated damage promotes the GAP activity of TBC1D15. Higher Rab7-GAP activity of TBC1D15 may be necessary to segregate the damaged lysosomal mass from vesicular traffic. Further structural and biophysical experimentation is required to understand whether and how LC3 binding of TBC1D15 is influenced by its GAP activity.

### TBC1D15 is needed to propagate lysosomal regeneration flux

Having confirmed the recruitment of TBC1D15 to damaged lysosomes, we investigated its role in lysosomal regeneration using tfGal3 (**Fig 1B**). HeLa cells were treated with LLOMe for 2 h, followed by washout in LLOMe-free medium for the same duration. As expected, the number of red Gal3 puncta increased after recovery for 2 h, indicating the regeneration of lysosomes after damage (**Fig 3A, 3B**). Interestingly, TBC1D15 colocalized with yellow Gal3 puncta representing damaged lysosomes, but not with red Gal3 puncta representing recovered, acidic lysosomes (**Fig 3A, 3B**). Next, we depleted endogenous TBC1D15 in HeLa cells using the corresponding siRNA and then reconstituted with wild-type, LIRmut or GAPmut TBC1D15 before repeating the flux assay. The absence of TBC1D15 completely blocked the lysosomal regeneration flux. The phenotype could be rescued with the wild-type protein, but not with either of the mutants. These findings confirm that TBC1D15 is required for the regeneration of lysosomes after damage, and that the LIR and GAP domains are both necessary for this function (**Fig 3C**).

**Figure 3:**
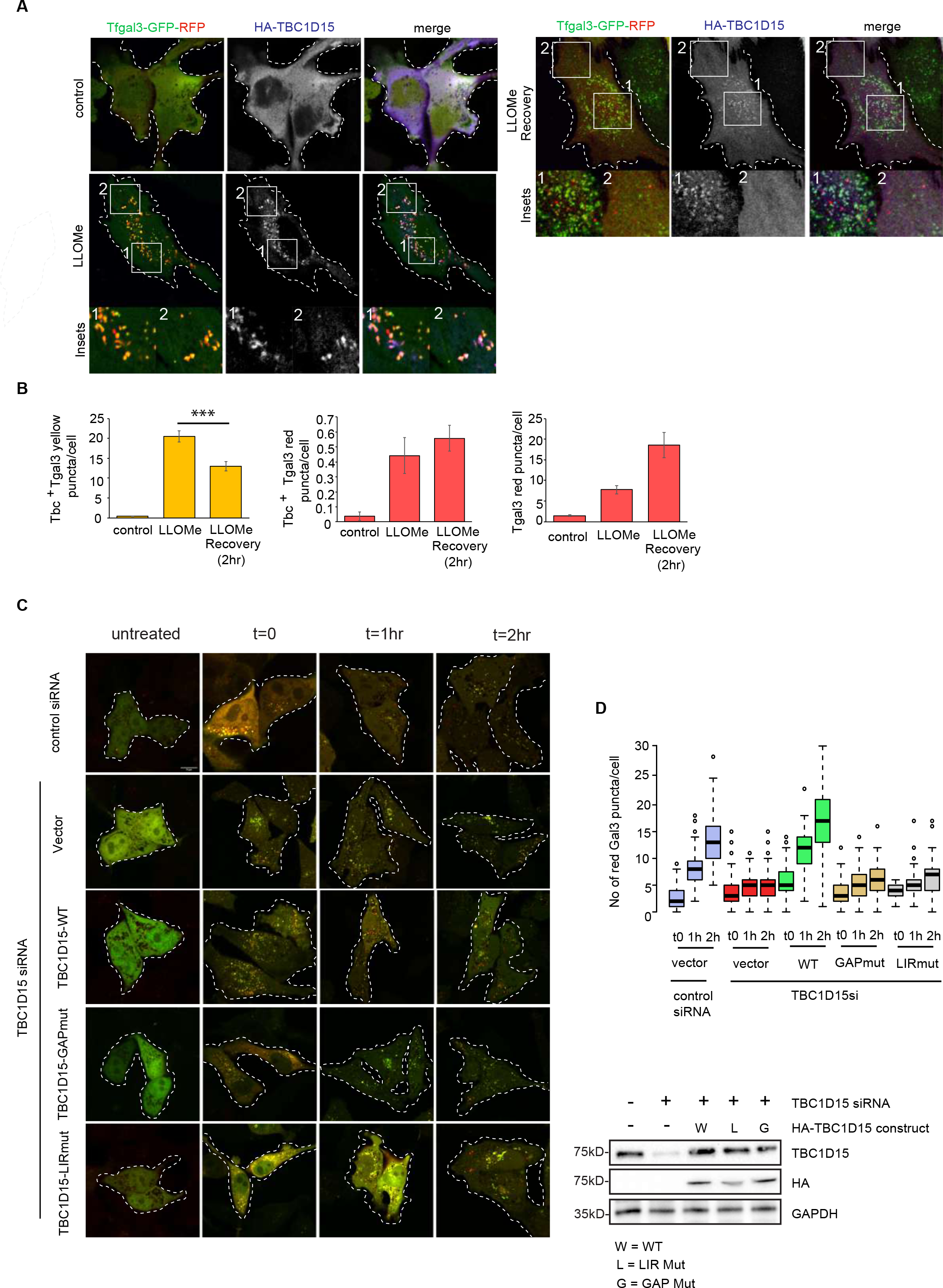
TBC1D15 is necessary for lysosomal regeneration. a) HeLa cells with stable expression of the dual-tagged tfGal3 reporter and HA-TBC1D15 were treated with 1 mM LLOMe for 2 h followed by washout (LLOMe recovery). Cells were then fixed and stained with an antibody specific for HA for confocal imaging. Red gal3 puncta correspond to regenerated lysosomes. b) Images in panel (a) were analysed using FIJI to count the numbers of TBC1D15 ^+^ yellow gal3 puncta, TBC1D15^+^ red gal3 puncta, and the total number of red gal3 puncta/cell to follow damage and recovery events. Data are means ± SEM of at least 50 cells counted from three independent experiments (***p≤0.001). c) Endogenous TBC1D15 was silenced with siRNA for 48 h in HeLa cells with stable expression of tfGal3. Wild-type (WT) and mutant TBC1D15 constructs were then transfected into the cells for reconstitution. The cells were then treated with 1 mM LLOMe for 2 h followed by LLOMe washout for 1 or 2 h. Expression levels of TBC1D15 were determined by immunoblotting of lysates made from cells used for imaging. d) The number of red gal3 puncta was counted per cell for each condition in the experiment in panel (c). Data are means ± SEM of 53 cells counted for each condition.

### Autophagy is important forTBC1D15 mediated lysosomal regeneration

Since LC3 binding to TBC1D15 was necessary for its lysosomal localisation and for the propagation of lysosomal regeneration flux in LLOMe treated cells, we explored the role of autophagy in TBC1D15 mediated lysosomal regeneration. To check this, we treated cells with siRNA targeting different molecules of autophagy: ATG13 (a part of the pre-initiation complex), ATG3 and ATG12 (important for lipidation of ATG8 proteins), and Rubicon (essential for the non-canonical autophagy type LC3 associated phagocytosis (LAP)). Depletion of ATGs (ATG13, ATG3, ATG12) led to a significant decrease in TBC1D15 recruitment to damaged lysosomal membranes; while the depletion of Rubicon did not have any effect on TBC1D15 recruitment (**Fig. S3A**). The siRNA-mediated depletion of ATG16L also reduced TBC1D15 recruitment to lysosomes. Reconstitution with both WT and the CASM deficient mutant of ATG16L (K490A) (Fletcher et al., 2018) restored the recruitment of TBC1D15 to damaged membranes (**Fig. S3B**).

Further, we treated cells expressing LC3-BFP with the following pharmacological inhibitors of autophagy: the PI3K inhibitor Wortmannin, the ULK1/2 inhibitor MRT68921 and the Vps34 inhibitor SAR405, in presence of LLOMe and monitored the formation of TBC1D15 puncta. Treatment with all the three drugs conspicuously reduced the recruitment of TBC1D15 to lysosomes (**Fig. S4A and B**). Also, HA-TBC1D15 from similarly treated cells did not interact with LC3B (**Fig. S4C**). Lysosomal regeneration flux measured in cells stably expressing tfGal3 also compromised upon pharmacological inhibition of autophagy (**Fig S4D and E**). Collectively, these results suggest that the autophagy (classical macroautophagy) pathway is important for TBC1D15 mediated lysosomal regeneration but non-canonical autophagy forms involving lipidation of ATG8 on single membranes (like LAP and CASM) are not required for the process.

### Proximity labelling of TBC1D15 confirms its role in lysosomal regeneration

To determine how TBC1D15 facilitates lysosomal regeneration, it was important to understand the molecular mechanism underlying the restoration of lysosomes after damage. We therefore applied a proximity labelling method in which inducible Flp-in Trex HeLa cells expressing APEX2-TBC1D15 under doxycycline treatment were treated with LLOMe or left untreated, followed by labelling with biotin using a 1-min pulse of 1 mM H_2_O_2_. The biotin-labelled proteome was pulled down using streptavidin beads and analysed by MS (**Fig 4A and B**). We compared LLOMe-treated samples to the untreated control and filtered out significant hits after false discovery rate (FDR) correction using Perseus software. Gene Ontology (GO) analysis of significantly enriched proteins identified the lysosomal membrane proteins SCARB2, TMEM9 and TMEM192, the lysosomal V-ATPase (ATP6V0-V1) complex, and hydrolytic enzymes such as β-hexosaminidase (HEXA, HEXB), α-glucosidase (GAA), and γ-glutamyl hydrolase (GGH). Autophagy-related proteins such as GABARAPL2, FIP200 (RB1CC1) and ATG3 were also enriched by LLOMe treatment. We also detected enriched proteins involved in lipid metabolism, including PLCH1, PLCB3, PLPP6 and GPD2 (**Fig 4C**).

**Figure 4:**
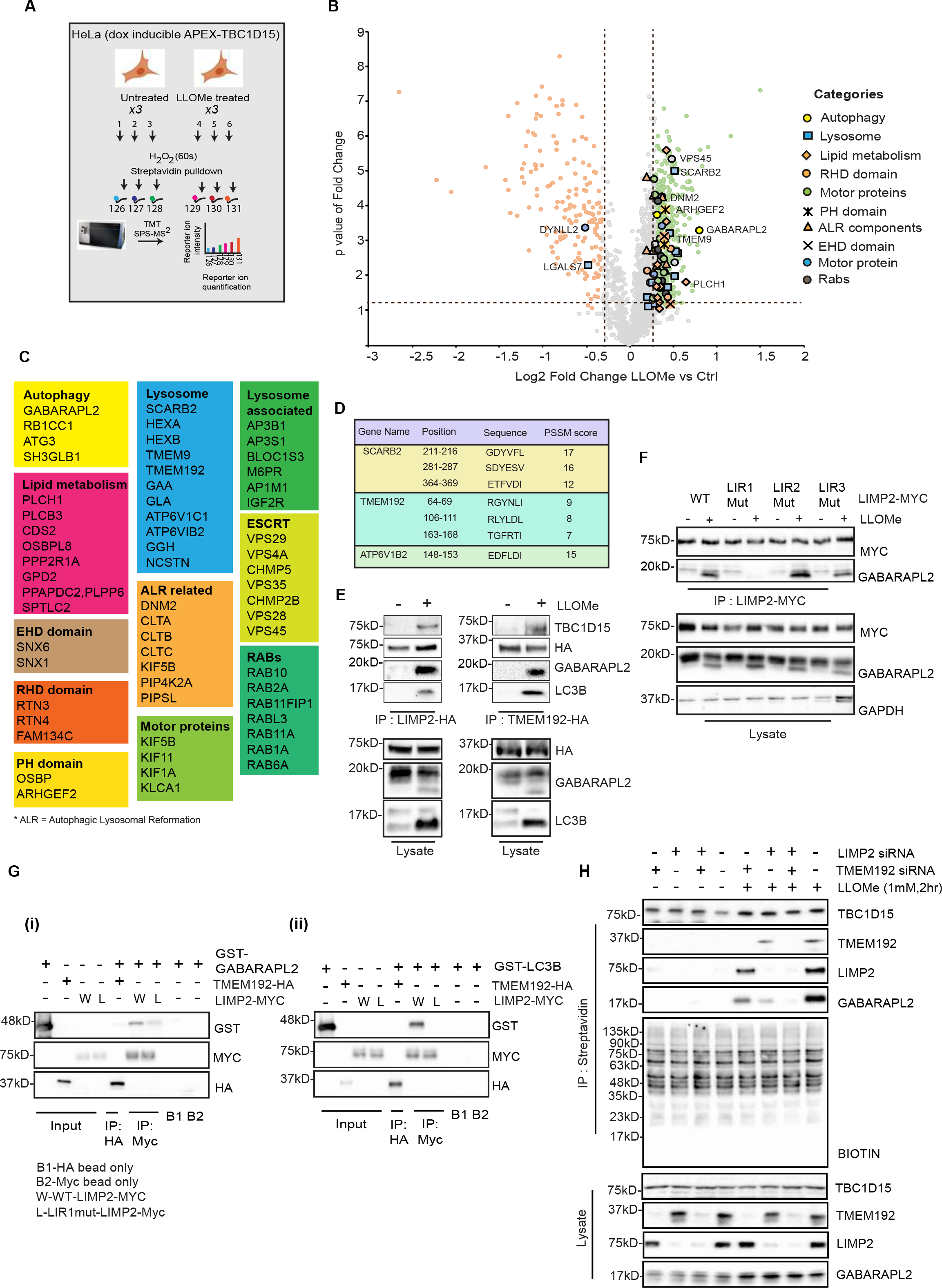
Proximity labelling of TBC1D15 identifies LC3B-binding proteins and ALR machinery near damaged lysosomal membranes. a) Schematic of the APEX2-TBC1D15 proximity labelling assay. b) Volcano plot showing biotinylated proteins enriched by LLOMe treatment. c) Pathway analysis of proteins enriched by LLOMe treatment from the experiment in panel (b) d) Lysosomal membrane proteins, significantly enriched in panel (b), were scanned for potential LIR motifs using iLIR database. Putative LIR motifs are listed. e) HEK 293T cells were transfected with HA-tagged LIMP2 and TMEM192 constructs, then treated with 1mM LLOMe for 2 h to induce lysosomal damage. The extent of interaction with ATG8 proteins and TBC1D15 was assessed by HA immunoprecipitation followed by western blotting. f) Putative LIR motifs for LIMP2 as described in panel (d) were mutated by site directed mutagenesis. MYC-tagged LIMP2 constructs were transfected in HEK 293T cells and then were treated with 1mM LLOMe for 2 h or left untreated. The interaction with GABARAPL2 and LIMP2 was assessed by MYC immunoprecipitation and western blotting. g) Wild type TMEM192 and wild type or LIR1 mut LIMP2 variants were purified from HEK293T cells and incubated with pure GST-GABARAPL2 and GST-LC3B and extent of binding were assessed by MYC and HA immunoprecipitation followed by western blotting. Efficiency of binding with pure GABARAPL2 and LC3B is shown in panel (g-i) and (g-ii) respectively. h) HeLa cells expressing doxycycline-inducible APEX2-TBC1D15 were treated with siRNA to deplete LIMP2 and TMEM192 either individually or in combination. TBC1D15 expression was then induced with doxycycline, followed by LLOMe treatment and proximity labelling with biotin/H_2_O_2_. Biotinylated proteins were pulled down using streptavidin-agarose and mentioned proteins were checked by western blotting using specific antibodies.

### LC3-binding proteins on lysosomes interact with TBC1D15

We next sought LC3 binding proteins on damaged lysosomes, given that TBC1D15 is not responsible for this role (**Fig 2H**). The lysosomal proteins found in the proximity of TBC1D15 in LLOMe-treated cells were screened against the iLIR database, revealing three proteins with putative LIR motifs: ATP6V1B2, TMEM192 and SCARB2 (LIMP2) (**Fig 4D**). Immunoprecipitation of HA-tagged TMEM192 and LIMP2 from LLOMe-treated cells showed that both proteins interacted with GABARAPL2 and LC3B, whereas endogenous ATP6V1B2 did not interact with ATG8 proteins. Interestingly, TMEM192 and LIMP2 also interacted with endogenous TBC1D15 in LLOMe-treated cells (**Fig 4E, S5A**). Mutation of the three putative LIR motifs of LIMP2 demonstrated that LIR1 (amino acids 211-216, GDYVFL) is the active LIR motif that mediates its interaction with GABARAPL2 in LLOMe treated cells (**Fig. 4F**). Mutation of the predicted LIR motifs of TMEM192 did not affect its interaction with GABARAPL2 (**Fig. S5B**). Next, purified WT, the LIR mutant (LIMP2[Y213AF215AL216A], LIR1mut) of LIMP2-Myc and WT TMEM192-HA from HEK293T cells were tested for their interaction with GST tagged ATG8 proteins (GABARAPL2 and LC3B) purified from *E. coli.* Interaction assays using pure proteins showed that LIMP2-Myc binds to GST tagged LC3B and GABARAPL2 using its LIR motif (GDYVFL) since mutation of this sequence caused a significant reduction in protein-protein interaction. TMEM192 does not interact with LC3B or GABARAPL2 (**Fig 4G** **(i) and (ii)**). This suggests that LIMP2 directly interacts with ATG8 proteins using its LIR motif in LLOMe treated cells. TMEM192 may be a part of a larger complex of proteins which bind ATG8s during lysosomal damage. Next, we depleted endogenous LIMP2 and TMEM192 by the siRNA treatment of HeLa cells expressing APEX2-TBC1D15 followed by LLOMe treatment and biotin labelling. The biotin-labelled proteome in these cells was then mixed with streptavidin-agarose and probed with antibodies against these proteins, biotin, and GABARAPL2. The siRNA-mediated knockdown of LIMP2 and TMEM192 reduced the amount of GABARAPL2 biotinylation, suggesting less interaction with TBC1D15 (**Fig 4H**). The binding of ATG8 proteins to LIMP2 and TMEM192 may therefore be necessary for the recruitment of TBC1D15 to damaged lysosomes. To test this hypothesis, we depleted endogenous LIMP2 and TMEM192 with siRNA for 48 h and checked the recruitment of TBC1D15 to damaged lysosomes by microscopy. The knockdown of TMEM192 had no significant impact on the number of TBC1D15^+^ lysosomes; knockdown of LIMP2 causes a small but statistically significant reduction of TBC1D15 recruitment to lysosomes, while cells depleted for both LIMP2 and TMEM192 had significantly fewer TBC1D15 puncta after LLOMe treatment, and these colocalized with lysosomes (**Fig S5C-E**). Depletion of these lysosomal proteins also reduced lysosomal regeneration flux when measured with the tfGal3 reporter assay (**Fig. S5F and G**).

The lysosomal calcium channel TRPML1 binds ATG8 proteins following the treatment of cells with its agonist MLSA1, which releases Ca^2+^ from lysosomes (Nakamura et al., 2020). Given that the binding of ATG8 proteins to LIMP2 triggered TBC1D15 recruitment to lysosomes, we checked whether TBC1D15 was also recruited to lysosomes after MLSA1 treatment. TBC1D15 puncta colocalized with lysosomes when cells were treated with 50 µM MLSA1 for 1 h (**Fig S5H**). Under this condition, TRPML1-APEX2 is in proximity to LC3, GABARAPL2 and TBC1D15 (**Fig S5I**). Taken together, these results suggest that TBC1D15 is indeed a part of the lysosomal membrane damage response and is recruited to lysosomes under conditions where the ATG8 proteins bind to LIMP2.

### TBC1D15 interacts with the lysosomal regeneration machinery

Analysis of the significant hits from the APEX2-TBC1D15 proximity labelling assay using Metascape for pathway and process enrichment (Zhou et al., 2019) identified vesicle-mediated transport, and a cluster of ALR proteins (including DNM2, KIF5B and CLTC) (**Fig S6A**), as the top hits (Yu et al., 2010). We hypothesized that ALR components may contribute to the regeneration of active lysosomes from damaged membranes following LLOMe treatment. Our proximity labelling analysis of APEX2-TBC1D15 suggested that TBC1D15 is in proximity to ALR proteins in LLOMe-treated cells (**Fig 5A**). Like in our LAMP1-APEX2 proximity labelling experiment (Fig 2B), LC3 was noted to be in the proximity of TBC1D15 in this assay.

**Figure 5:**
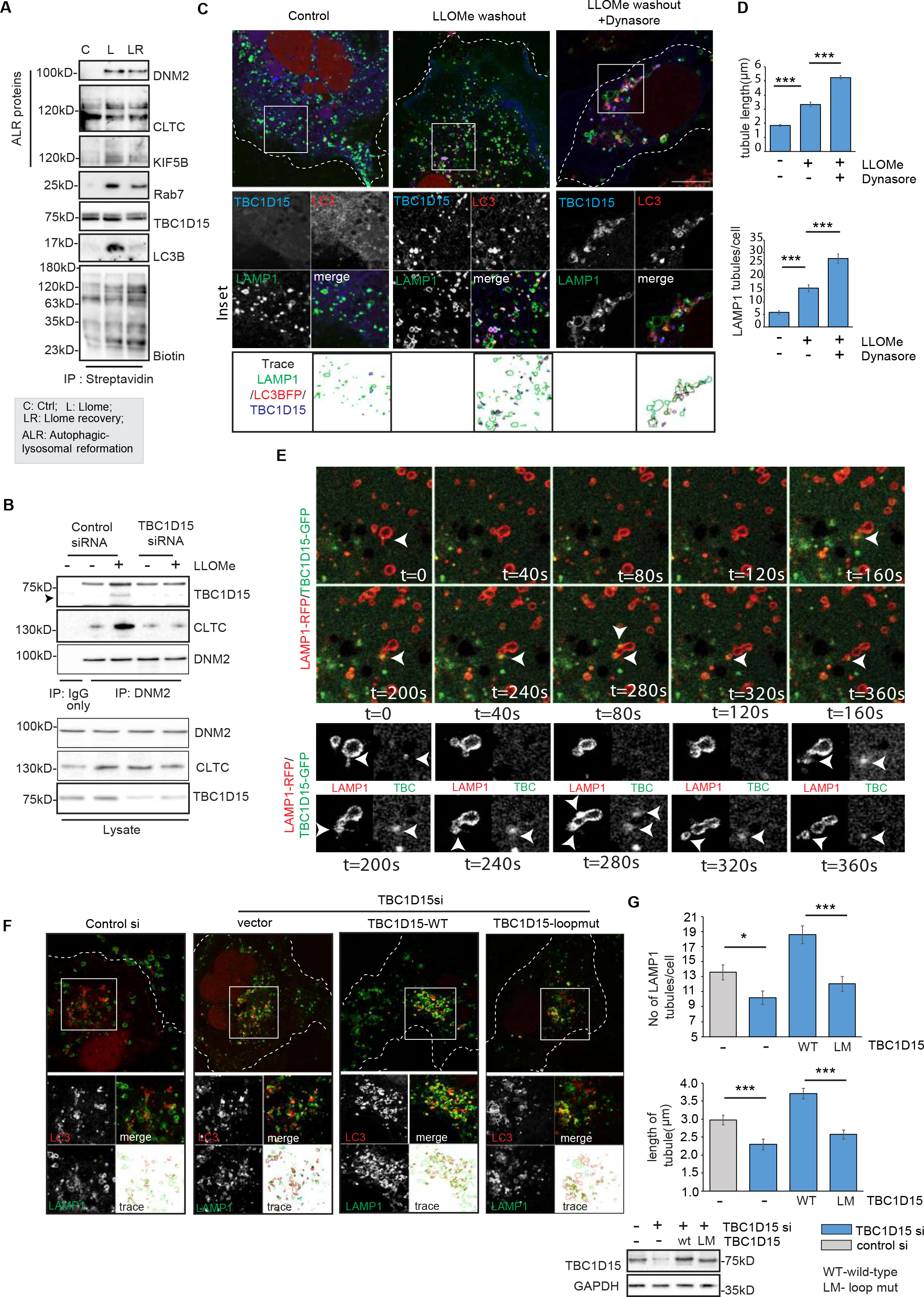
Lysosomes regenerate by TBC1D15-dependent tubulation. a) Cells expressing doxycycline-inducible APEX2-TBC1D15 were treated as indicated and exposed to biotin/H_2_O_2_ before a proximity labelling assay. Following streptavidin pulldown, samples representing different conditions were analysed by western blotting and the proteins of interest (DNM2, CLTC, KIF5B, Rab7, TBC1D15 and LC3B) were detected using specific antibodies. C = control; L = treatment with 1 mM LLOMe for 2 h; LR = LLOMe treatment followed by 2 h recovery; ALR = autophagic lysosomal reformation. b) Endogenous DNM2 was immunoprecipitated from TBC1D15-depleted cells (untreated or treated with 1 mM LLOMe, 2 h) and the interactions of DNM2 with CLTC and TBC1D15 were assessed by western blotting. c) Cells expressing LC3B-BFP were treated with 1mM LLOMe for 2hr followed by recovery in LLOMe free medium with or without 20 µM Dynasore for 2 h, and were fixed and stained for TBC1D15 and LAMP1. These samples were then imaged by airy-scan confocal microscopy. Images were analysed in FIJI to obtain the trace of LAMP1 membranes (green), LC3BFP (red) and TBC1D15 (blue) d) Cells from experiment in panel (c) were analysed with FIJI to measure the length and number of lysosomal tubules. LAMP1 structures greater than 2.5 µm in length were defined as a lysosomal tubule when counting number of tubules per cell. Data are means ± SEM of 60 cells from three independent experiments (***p≤0.001). e) Time-lapse imaging of cells expressing LAMP-RFP and TBC1D15-GFP treated with 1mM LLOMe for 2 h followed by LLOMe recovery for 1 h. Arrows indicate position of TBC1D15 on the LAMP1 tubule. f) HeLa cells were treated with TBC1D15-specific siRNA to deplete the endogenous pool followed by reconstitution with either full-length TBC1D15 or the loop mutant. These cells were then treated with 1mM LLOMe for 2 h and were allowed to recover in LLOMe free medium for 2 h. Cells were then fixed and stained for TBC1D15 and LAMP1 and lysosomal tubulation events were observed by airyscan confocal microscopy. Images were analysed in FIJI to obtain the trace of LAMP1 membranes (green), LC3BFP (red) and TBC1D15 (blue) g) Cell counts corresponding to the images in panel (f). Data are means ± SEM of 50 cells counted for all conditions to compare the number and length of LAMP1 tubules (***p≤ 0.001, *0.01≤p < 0.05).

### The TBC1D15 interactome also reveals the assembly of ALR components on damaged lysosomal membranes

The interactome of HA-TBC1D15 revealed that DNM2 and KIF22 may interact directly with TBC1D15 in LLOMe-treated cells (**Fig S6B**). We also captured ALR proteins following the enrichment of damaged lysosomal membranes. In this case, cells expressing TBC1D15-GFP, LAMP1-RFP and LC3-BFP were treated with LLOMe and permeabilised with 0.05% digitonin to reduce cytosol and enrich for cellular membranes. Confocal microscopy after digitonin treatment revealed the extensive colocalization of TBC1D15, LAMP1 and LC3 in the LLOMe-treated samples and this signal did not diminish even after digitonin treatment (**Fig S6C**). GFP pulldown from TBC1D15-GFP expressing cell lysates after digitonin treatment, followed by TBC1D15-GFP interactome analysis, revealed the presence of LC3B and the ALR components CLTC and KIF5B in the interactome during LLOMe treatment (**Fig S6D**).

### The TBC1D15 N-terminal domain is a scaffold for the assembly of ALR components

DNM2 was found in most of our TBC1D15 interactome datasets, so next we tested whether DNM2 interacts directly with TBC1D15. The C-terminal portion of TBC1D15 contains the catalytic GAP domain and LIR motifs, and has already been structurally characterized (Chen et al., 2017). However, little is known about the structure of the N-terminal portion, so we analyzed the complete protein using the structure from the AlphaFold database (Jumper et al., 2021). The predicted structure of TBC1D15 featured the N-terminal domain and C-terminal domain connected by a linker (**Fig S7A**). To identify potential interaction motifs, we expressed full-length GST-tagged TBC1D15 in *Escherichia coli*, and versions without the N-terminus (ΔNter, TBC1D15 amino acids 301–697) and without the C-terminus (ΔCter, TBC1D15 amino acids 1–399). We then utilized the purified proteins for GST pulldown assays using the lysates of LLOMe-treated cells. The N-terminal portion of TBC1D15 bound to DNM2 and CLTC (**Fig S7B**) and the testing of mutations revealed that the first 190 amino acids were most important for the interaction with DNM2, especially residues 102–183 (**Fig S7C**). The AlphaFold model predicted the presence of a loop spanning residues 57–124, which we considered a likely structure to interact with DNM2 (**Fig S7D**). We therefore generated a mutant of the full-length protein in which this loop was completely deleted (TBC1D15Δ[57-124], loop mutant). The interaction between TBC1D15 and DNM2 was weaker when the loop region was deleted (**Fig S7E**). We concluded that the C-terminal region of TBC1D15 is responsible for LC3 binding and its GAP activity while the N-terminal region interacts with DNM2. Endogenous DNM2 interacted with TBC1D15 and CLTC in LLOMe-treated cells but the interaction between DNM2 and CLTC was weaker following the depletion of TBC1D15 (**Fig. 5B**), suggesting that TBC1D15 acts as a scaffold to assemble the ALR molecular machinery. Whether the interaction between TBC1D15 and ALR components are direct or whether TBC1D15 is a part of a multi-protein complex that also harbours ALR proteins require further testing.

### The interaction between TBC1D15 and DNM2 is important for lysosomal regeneration

To establish the role of TBC1D15 and DNM2 in lysosomal regeneration, we repeated the RFP-GFP-Gal3 flux assay in the presence of dynasore, which inhibits the DNM2-mediated scission of lysosomal tubules. Dynasore treatment or depletion of DNM2 with siRNA significantly reduced the number of red Gal3 puncta after 2 h of LLOMe washout (**Fig S7F and G**). This was also the case in cells treated with siRNA to deplete endogenous TBC1D15. The effect could be rescued by expressing wild-type TBC1D15 but not the loop mutant, confirming the importance of the TBC1D15-DNM2 interaction in the process of lysosomal regeneration (**Fig S7H**).

### Lysosomal tubules form proto-lysosomes in response to LLOMe-mediated damage

Lysosomal regeneration involves the formation of tubules from LC3^+^ structures, which are subsequently cleaved by DNM2 to yield proto-lysosomes (Yu et al., 2010; Khundadze et al., 2021). High resolution (STED) imaging of endogenous LAMP1 in untreated, LLOMe treated, and during LLOMe washout conditions was performed: in untreated cells, LAMP1 was present on lysosomes. In LLOMe treated cells, the LAMP1 staining was dispersed indicating loss of lysosomal membrane integrity. Upon LLOMe washout, short LAMP1 positive membranes, longer curved tubules resembling open ring, and larger closed vesicles were evident (**Figure S8A and C**). Depletion of TBC1D15 caused a decrease in formation of tubules while treatment with DNM2 siRNA increased the length of these LAMP1 tubules (**Figure S8B and C**). To determine how LC3B and TBC1D15 were distributed on the LAMP1 positive structures, we treated HeLa cells expressing LAMP1-RFP, TBC1D15-GFP and LC3-BFP with LLOMe, followed by LLOMe washout and imaged them by airy-scan confocal microscopy. LAMP1-GFP tubules were observed, often curved in the form of open (unsealed) rings. TBC1D15 and LC3B were closely associated with LAMP1 structures. Treatment with dynasore led to formation of longer tubules which often appeared like a tangled mass of membranes at airy-scan resolution; indicating Dynamin2 activity is important for scission of these tubules (**Fig 5C, 5D, S9A and S9B, supplementary videos 1, 2**). Time-lapse imaging of cells expressing LAMP1-RFP and TBC1D15-GFP after LLOMe treatment followed by 2 h of LLOMe washout showed the presence of large LAMP1 positive closed rings. TBC1D15 was present on certain regions of these LAMP1 rings. Small membrane buds protrude from these LAMP1 rings, are pulled and cleaved to form small LAMP1 positive vesicles (proto-lysosomes) in an ALR-like manner (**Fig 5E, S9C, supplementary videos 3, 4**). TBC1D15 depletion reduced the number and length of LAMP1 tubules in the cells. This phenotype could be rescued by reconstitution with wild-type TBC1D15 but not the loop mutant (**Fig. 5F, 5G**). Tubules observed by airy scan confocal imaging in Fig. 5C and Fig. 5G were LC3^+^ or LAMP^+^ or have both markers suggesting they are a mixture of lysosomal and autolysosomal membranes. These experiments support our biochemical findings and confirm that, following extensive damage, the impaired lysosomal mass undergoes a regeneration process that requires TBC1D15 and the ALR machinery to create proto-lysosomes.

### TBC1D15 regulates lysosomal regeneration in response to damage caused by calcium oxalate

Finally, we tested the role of TBC1D15 in lysosomal regeneration using a model of crystal nephropathy. Previous studies have shown that lysosomal damage marked by GFP–Gal3 puncta occurs in immortalized kidney proximal tubular epithelial cells (PTECs) exposed to calcium oxalate crystals, causes the lipidation of ATG8 on the lysosomal membrane and the nuclear translocation of TFEB (Nakamura et al., 2020). In our system, oxalate-induced damage led to the formation of Gal3-GFP puncta (**Fig 6A**). HA-TBC1D15 was recruited to damaged lysosomes in cells treated with calcium oxalate (**Fig 6B**). Oxalate-induced lysosomal damage also led to the formation of LAMP1-GFP tubules characteristic of lysosomal regeneration. Treatment with dynasore increased the number of tubules, showing that DNM2 is needed to regulate the cleavage of proto-lysosomal tubules (**Fig 6C**). The depletion of TBC1D15 led to a significant decrease in tubule number and length (**Fig 6D**). Our PTEC/calcium oxalate model therefore confirmed that lysosomal regeneration takes place during oxalate nephropathy and is dependent on TBC1D15 and DNM2.

**Figure 6:**
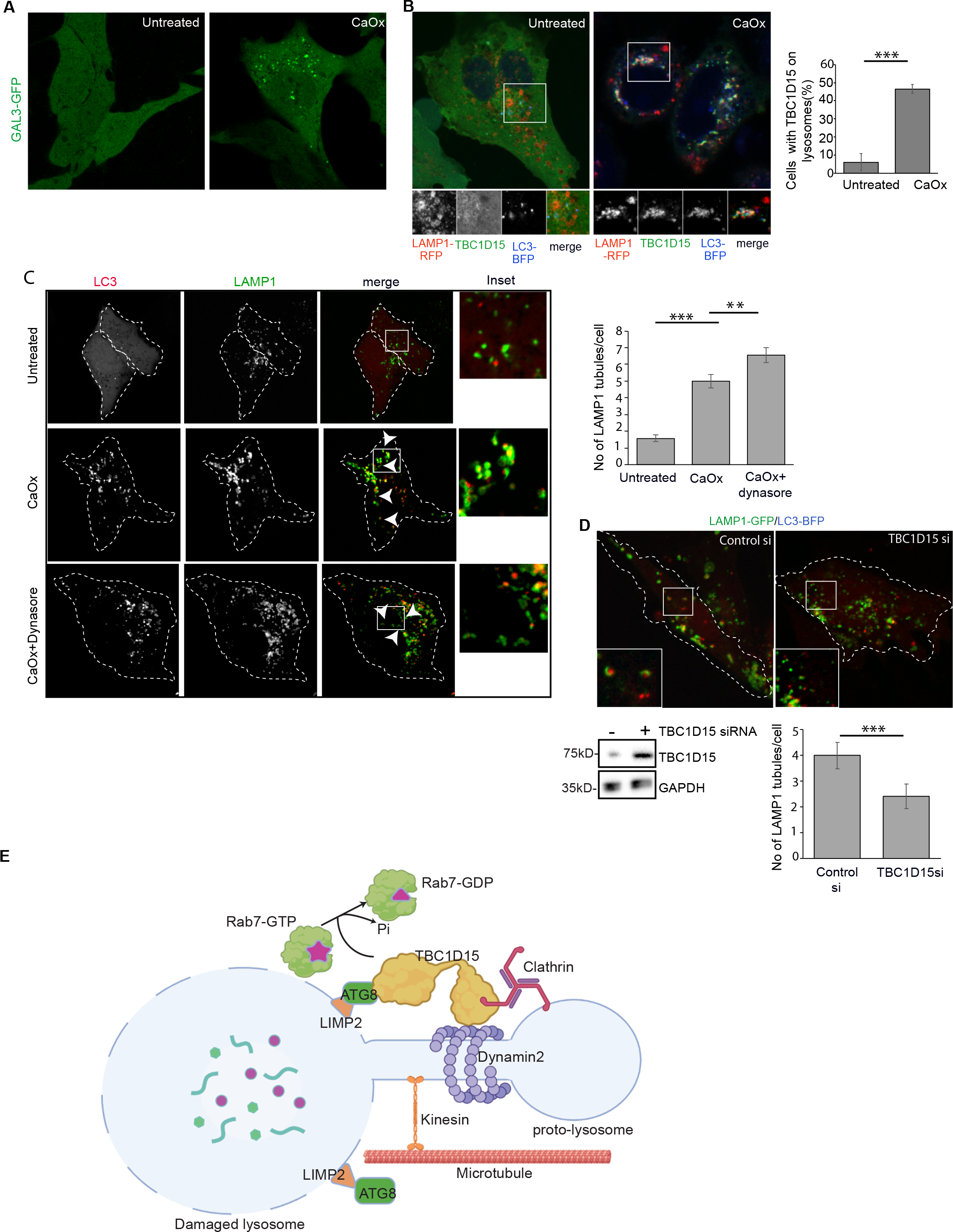
TBC1D15 regulates lysosomal tubulation in PTEC cells treated with calcium oxalate. a) PTEC cells expressing GFP-GAL3 were treated with 100 μ/ml calcium oxalate (CaOx) for 2 h to induce lysosomal damage, and the extent of damage was determined by the analysis of GFP-Gal3 puncta. b) PTEC cells were transfected with LAMP1-RFP and LC3-BFP, and the colocalization of TBC1D15 with LAMP1 was assessed by confocal microscopy based on the staining of endogenous TBC1D15. Data are means ± SEM of at least 100 cells from three independent experiments (***p≤0.001). c) PTEC cells expressing LAMP1-RFP and LC3-BFP were treated with 100 g/ml CaOx or 100 g/ml CaOx plus 20 µM Dynasore to check for lysosomal tubulation events. Data are means ± SEM of 70 cells imaged to determine the number of lysosomal tubules under different conditions (***p≤0.001, **0.001 < p≤0.01). d) PTEC cells expressing LAMP1-GFP and LC3-BFP were treated with either control siRNA or siRNA specific for TBC1D15 and then with 1 mM LLOMe to determine the number of LAMP1 tubulation events. Data are means ± SEM of 50 cells counted to compare the two conditions (***p≤0.001). Knockdown efficiency of TBC1D15 was checked by western blotting. e) Summary of the response to acute lysosome damage. TBC1D15 is recruited to the damaged membrane mass by interacting with ATG8 proteins bound to lysosomal membrane proteins. TBC1D15 hydrolyses Rab7-GTP to segregate this damaged mass from vesicular traffic. The LIR and GAP functions are localized to the C-terminal portion of TBC1D15, whereas the N-terminal portion acts as a scaffold to recruit ALR machinery at damaged foci, which potentiates lysosomal regeneration.

## Discussion

We have identified a new pathway of lysosomal membrane regeneration that rapidly generates functional lysosomes independently of lysosomal biogenesis. This process requires TBC1D15, DNM2 and ATG8 binding to the lysosomal membrane protein LIMP2. Lysosomal regeneration shares common features with ALR. Both processes require ATG8 proteins to initiate the formation of lysosomal tubules, which are elongated by pulling along microtubules using kinesins, and then cleaved by DNM2 to generate new lysosomes. Analysis of a TBC1D15 structural model followed by mutational analysis showed that the TBC1D15 C-terminal domain binds to ATG8 proteins while the N-terminal domain interacts with DNM2. TBC1D15 is anchored to damaged lysosomes by interacting with ATG8 proteins bound to lysosomal membrane proteins, and acts as a scaffold to assemble the regeneration machinery that is needed to produce new lysosomes (**Fig 6E**).

The recruitment of TBC1D15 to lysosomes is dependent on its LIR motif. We identified LIMP2 and TMEM192 as potential lysophagy receptors in our proximity-labelling experiments. However, experiments with pure proteins showed that LIMP2 binds to ATG8 proteins utilising its LIR motif while TMEM192 is not a direct interactor. Depletion of LIMP2 (but not TMEM192) caused a reduction in lysosomal regeneration flux but the concurrent depletion of both these genes led to a greater decrease in lysosomal regeneration. This suggests that though TMEM192 is not a lysophagy receptor, it may be a part of the multi-protein complex which binds ATG8 proteins in LLOMe treated cells. LIMP2 binding to ATG8 proteins may be responsible for the specificity of TBC1D15 to damaged lysosomal membranes since depletion of LIMP2 significantly reduces lysosomal recruitment of TBC1D15 even though LC3B is still recruited to lysosomes.

The lysosomal calcium channel TRPML1 was recently shown to bind lipidated LC3 following treatment with its agonist MLSA-1 (Nakamura et al., 2020). Our experiments with TRPML1-APEX2 showed that MLSA1, which brings ATG8 proteins in the proximity of TRPML1 also has TBC1D15 in the vicinity. Further studies on the molecular and structural basis of these observations are needed to achieve better insight into the specificity of TBC1D15 recruitment and its function in this scenario.

Formation of autophagosomal membranes is also essential for the TBC1D15 dependent pathway of lysosomal regeneration. Pharmacological inhibition of autophagosome formation or siRNA mediated depletion of autophagy genes led to a decrease in TBC1D15 mediated lysosomal regeneration flux. Lipidated LC3 present on autophagosomal membranes binds to lysophagy receptors (like LIMP2) on the damaged lysosome. In presence of TBC1D15, damaged lysosomal membranes form elongated tubules, open or closed LAMP1 rings which are quite distinct in morphology when compared to the tubules formed during ALR (Yu et al., 2010, Khundadze et al., 2021). TBC1D15 is important for formation of these LAMP1 positive structures, in presence of TBC1D15, membrane buds are formed which are pulled along microtubules and cleaved by Dynamin2 to form new LAMP1+ proto-lysosomes in a process which closely resembles ALR. These protolysosomes are acidic, but have sub-optimal hydrolase activity when compared to that of mature lysosomes in untreated cells. To avoid ambiguity, we have called this process lysosomal regeneration since the process is similar but not identical to ALR.

TBC1D15-mediated lysosomal regeneration requires the ALR machinery to form new lysosomes, but other proteins may also be involved. The biotinylated TBC1D15 interactome revealed several groups of proteins that may regulate the process, including ESCRT proteins that are well-known for their role in membrane budding and scission. Many other interacting proteins featured lipid-binding domains such as the PH domain (OSBP, ARHGEF2, KIF1A and OSBPL8), BAR domain (SNX6 and SH3GLB1), C2 domain (PLCH1, RABL1FIP1, PLCB3 and ESYT2), and ENTH domain (HIP1 and PICALM), suggesting that the regeneration of damaged membranes involves the complex remodelling of lipids, although the exact processes remain unknown.

Finally, it is important to understand the physiological context in which lysosomal regeneration may come into play. The lysosomal damage response is sensitive to the duration and extent of damage, with minor membrane perturbations triggering the calcium-dependent recruitment of ESCRT proteins for membrane repair (Skowra et al., 2018; Radulovic et al., 2018), but greater damage triggering lysophagy, where lysosomal membrane proteins bind to ATG8 proteins and recruit selective autophagy receptors such as TAX1BP1, SQSTM1 and NDP52 (Koerver et al., 2019; Eapen et al., 2021). Autophagosomes with cargos of damaged organelles then fuse with healthy lysosomes. We found that acute damage reduces the proportion of functional (acidic) lysosomes far below that of damaged organelles, making it difficult to regenerate lysosomes through conventional autophagosome–lysosome fusion. This scenario may result in the induction of an instantaneous regeneration program, also requires autophagy proteins, but its contribution becomes significant when the damage is most severe (as seen after 2–3 h of treatment with LLOMe, when TBC1D15 distribution is almost completely lysosomal). This mechanism is probably used by the cell to generate a population of healthy lysosomes before TFEB/TFE3-mediated gene expression can initiate the program of lysosomal biogenesis. The treatment of PTECs with calcium oxalate also caused the recruitment of TBC1D15 to lysosomes and the formation of LAMP1 tubules, which are dependent on TBC1D15 and DNM2 for their formation and conversion to proto-lysosomes, respectively. The regeneration process may therefore contribute to the maintenance of lysosomal homeostasis in diseases such as crystal nephropathy.

## Materials and methods

### Cell culture

HeLa and HEK 293T cells were cultured in Dulbecco’s modified Eagle’s medium (DMEM, Gibco) containing 10% foetal bovine serum (FBS) and 1% penicillin–streptomycin. PTEC cells (a kind gift from Tamotsu Yoshimori, Department of Genetics, Osaka University, Japan) were grown in low-glucose DMEM (100 mg/dL glucose, Gibco) containing 10% FBS and 1% penicillin–streptomycin. Transfection with plasmid DNA and siRNA was carried out using Lipofectamine 2000 (Invitrogen) according to the manufacturer’s recommendations.

### Lysosomal regeneration flux assay using the stable tfGal3 reporter

For stable expression of tfGal3, HeLa cells were transfected with the tfGal3 construct (addgene number: 64149) and maintained in medium containing the selection marker Geneticin(800µg/ml) to obtain a stable expression of the protein. Lysosomal damage was induced by treating cells with 1mM LLOMe for 2 h, and recovery was performed by replacing LLOMe containing medium with LLOMe free medium after PBS wash for indicated time points. The cells were then fixed immediately with 4% paraformaldehyde, incubated with the appropriate primary antibodies, and red Gal3 puncta, detected by confocal microscopy as described above, were counted manually using FIJI. At least 100 cells from three independent experiments were used for statistical analysis.

### DQ-BSA assay for hydrolase activity

DQ-BSA green was purchased from ThermoFisher Scientific, cat no. D12050. HeLa cells were treated with siRNA to knockdown TFEB and then transfected with LAMP1-RFP to mark lysosomes. Cells were treated with 1mM LLOMe for 2 h and allowed to recover for different timepoints as mentioned. During the last 40 minutes of every sample, DQ-BSA was loaded into cells at a final conc. of 10µg/ml in prewarmed media. Cells were then imaged by confocal microscopy.

### Plasmid construction

For proximity labelling experiments, the TBC1D15 and LAMP1 coding sequences were tagged with the enzyme APEX2 and introduced into the doxycycline-inducible vector pcDNA5 by Gibson assembly. The dual-tag Galectin3 plasmid was acquired from Addgene (ref. 64149). The TBC1D15 coding sequence was subcloned into vectors pEGFPC1 and pGEX6P1 for expression in mammalian and bacterial cells, respectively. The LIRmut and GAPmut variants of TBC1D15 were generated by site-directed mutagenesis. The N-terminal (1–300) and C-terminal (400–697) segments, as well as nested N-terminal deletions ΔN1 (1–100), ΔN2 (1–182) and ΔN3 (1–300) and the loop mutant (57–124) were generated by standard restriction and ligation. The LIR mutations for LIMP2 and TMEM192 were introduced by site directed mutagenesis. The LIR mutants of LIMP2 are as follows; LIR1: GDYVFL mutated to GDAVAA, LIR2: SDYESV mutated to SDAEAA and LIR3: ETFVDI mutated to ETAVAA. The LIR mutants of TMEM192 are as follows; LIR1: RGYNLI mutated to RGANAA, LIR2: RLYLDL mutated to RLALAA and LIR3: TGFRTI mutated to TGAATI. The full-length and deletion constructs of TBC1D15 were subcloned into pGEX6P1 for bacterial expression. The HA-TBC1D15, LAMP1-GFP, LAMP-RFP, LC3-BFP, GST-GABARAPL2, GST-LC3B and GAL3-GFP constructs were sourced from our in-house plasmid collection. The siRNAs for TBC1D15 (sc-95660), TFEB (sc-38509), TFE3 (sc-38507), ATG3 (sc-72582), ATG12 (sc-72578), ATG13 (sc-97013), ATG16L (sc-72580), Rubicon (sc-78326), DNM2 (sc-35236), LIMP2 (sc-41546) and TMEM192 (sc-89327) were purchased from Santa Cruz Biotechnology.

### Drugs and treatments

Drugs used in this study are listed below:

| Name | Treatment conditions | Source |
| --- | --- | --- |
| LLOMe | 1mM, 2hr | Sigma, Cat ID: L7393 |
| Dynasore | 20 μM, 2hr | Sigma, Cat ID: D7693 |
| Wortmanin | 300 nM, 6hr | Sigma, Cat ID: W1628 |
| SAR405 | 10 μM, 6hr | Sigma, Cat ID: 5.33063 |
| MRT68921 | 10 μM, 6hr | Sigma, Cat ID: SML1644 |
| MLSA1 | 50 μM, 1hr | Sigma, Cat ID: SML0627 |
| Monensin | 100 μM, 1hr | Sigma, Cat ID: M5273 |
| Torin1 | 500 nM, 6hr | Sigma, Cat ID: 475991 |
| CCCP | 10 μM, 4hr | Sigma, Cat ID: C2759 |

### Protein purification

All GST-tagged proteins were purified from *E. coli* as previously described (Shin et al., 2020). Briefly, *E. coli* BL21(DE3) competent cells (New England Biolabs) were transformed with the corresponding plasmids and small cultures were grown in terrific broth (TB) overnight. These overnight cultures were used to seed larger-scale cultures that were incubated at 37 °C until the OD_600_ reached 0.6–0.8. Protein expression was then induced by adding 0.5 mM isopropyl-D-thiogalactopyranoside (IPTG) overnight at 18 °C. The bacteria were pelleted, resuspended in lysis buffer (50 mM Tris pH 7.5, 150 mM NaCl, 3 mM DTT) and lysed by sonication. The lysate was centrifuged (13,000 × g, 30 min, 4 °C) and the supernatant was passed through a 0.22-μ m before incubation with glutathione-Sepharose resin pre-equilibrated with lysis buffer for 2 h at 4 °C. Non-specific binders were removed by washing the resin three times in buffer containing 300 mM NaCl. Specific binding proteins were eluted (50 mM Tris.HCl pH 7.5, 150 mM NaCl, 15 mM glutathione) and exchanged to storage buffer (20 mM Tris-HCl pH 7.5, 100 mM NaCl).

### Antibodies

We used the following antibodies: TBC1D15 (cat. no. 121396; Abcam), GAPDH (cat. no. 2118; Cell Signaling Technology), GFP trap beads (cat. no. gta-100; ChromoTek), GFP (cat. no. sc-9996; Santa Cruz Biotechnology), LC3B (cat. no. 2775; Cell Signaling Technology), GABARAPL1+L2 (cat no. ab109364), DNM2 (cat. no. ab3457; Abcam), CLTC (cat. no. ab2731; Abcam), KIF5B (cat. no. MA1-19352; Thermo Fisher Scientific), Rab7 (cat. no. E907E; Cell Signaling Technology), TFEB (cat. no. 37785; Cell Signaling Technology), ATG16L (cat no. 8089, Cell Signaling Technology), TFE3 (cat no. HPA023881; Sigma), Biotin (cat. no. SC101339; Santa Cruz Biotechnology), HA (cat. no. 7392; Santa Cruz Biotechnology), Myc (cat no. sc-40; Santa cruz biotechnology), GST (cat. no. 2622; Cell Signaling Technology), LAMP1 (cat. no. 25630; Abcam), Ubiqiutin (cat. no. 3936; Cell Signaling Technology), ATP6V1B2 (cat. no. ab73404; Abcam), phospho-p70s6K Thr389 (cat no. 9205; Cell signalling technology), p70S6K (cat no. 9202; Cell signalling technology), ATG3 (cat no. 3415; Cell signaling technology), ATG12 (cat no. 2010; Cell signaling technology), ATG13 (cat no. 13468; Cell signaling technology), Rubicon (cat no. 68261; Cell signaling technology), TRPML1 (cat no. ab272608; Abcam) LIMP2 (cat no. 176317; Abcam. We also received aliquots of antibody raised in the lab as a kind gift from Paul Säftig, Institute of Biochemistry, Kiel University, Germany) and TMEM192 (cat. no. ab185545, Abcam). All primary antibodies were used at 1:1000 dilutions for western blotting and at 1:100 dilution for Immunofluorescence.

### Immunoprecipitation and western blotting

Cells were lysed in 50 mM Tris.HCl (pH 7.5) containing 150 mM NaCl and 1% Triton X-100 on a rotating wheel for 10 min at 4 °C. After pelleting the debris, the protein concentrations in the supernatant were measured using a BCA assay (Thermo Fisher Scientific). Immunoprecipitation was achieved by incubating cell lysates containing 1 mg of protein overnight with anti-HA agarose beads (Sigma-Aldrich), Myc-agarose beads (Santa cruz biotechnology: sc-40 AC) or GFP-Trap beads (Chromatek) in immunoprecipitation buffer (50 mM Tris.HCl pH 7.5, 150 mM NaCl, 0.5% Triton X-100). The beads were washed three times in wash buffer (50 mM Tris.HCl pH 7.5, 300 mM NaCl, 0.5% Triton X-100) to remove non-specific binders, and then boiled in 2× SDS sample buffer. We loaded 10% Tris-glycine gels with 20 µg of protein per lane. The samples were fractionated by SDS-PAGE at 150 V for 90 min before transfer to PVDF membranes at 300 mA for 2 h. The membranes were incubated in blocking buffer (5% bovine serum albumin) before western blotting with the appropriate antibodies.

### GST pulldown assay

Cells were lysed in 20 mM Tris.HCl (pH 7.5) containing 150 mM NaCl and 1% Triton X-100 and the lysates were incubated with 50μl GST beads for 2 h to reduce non-specific binding. The pre-cleared lysates were then pelleted and the protein concentration in the supernatant was measured as described above. We added 1 mg of the pre-cleared lysate to 20μl of glutathione-Sepharose resin containing 5μg of GST-tagged TBC1D15 and incubated the mixture overnight at 4 °C on a rotating wheel. Non-specific binders were removed by washing with 20 mM Tris.HCl pH 7.5 containing 300 mM NaCl and 1% Triton X-100 before SDS-PAGE and western blotting as described above.

### RILP pulldown assay

GST-tagged RILP was purified from *E. coli*, and 5μg of the pure GST-RILP protein was mixed with glutathione beads and 1 mg cell lysate in lysis buffer (50 mM Tris.HCl pH 7.5, 150 mM NaCl, 0.1% Triton X-100, protease inhibitor cocktail) overnight at 4 °C. Immunoprecipitated samples were washed (50 mM Tris.HCl pH 7.5, 300 mM NaCl, protease inhibitor cocktail), before SDS-PAGE and western blotting as described above, using an antibody against Rab7 to determine the proportion of GTP-bound Rab7. Whole cell lysates were blotted and probed with Rab7-specific antibodies to determine the level of total Rab7. Rab7-GTP levels were plotted by normalizing the signal intensities of the GST-RILP pulldown blots against total Rab7.

### Pure protein interaction assay

Myc tagged LIMP2 constructs (WT and LIR1 mut) and HA tagged TMEM192 construct (WT) were purified from HEK 293T cells by using Myc resin and HA resin respectively. These beads were washed three times with high salt wash buffer (50 mM Tris.HCl pH 7.5, 500 mM NaCl) to remove all cellular interactors. GST tagged ATG8 proteins were incubated with these enriched fractions of LIMP2 and TMEM192 for 1 h at room temperature followed by washing three times with wash buffer (50 mM Tris-HCl pH 7.5, 300 mM NaCl) to remove non-specific binding. These samples were then boiled with 2X SDS sample buffer and subjected to western blotting.

### Immunocytochemistry

HeLa cells growing on glass coverslips were fixed with 4% paraformaldehyde for 10 min at room temperature, permeabilised and blocked in phosphate-buffered saline (PBS) containing 0.1% saponin and 5% FBS for 1 h at room temperature, and incubated overnight at 4 °C with the appropriate primary antibody. Alexafluor-tagged secondary antibodies were used for visualization by fluorescence imaging.

### Confocal microscopy and image analysis

Confocal images were captured using our in-house Zeiss LSM780 microscope system fitted with a 63× 1.4 NA oil-immersion objective and argon and helium–neon lasers for the excitation of GFP and RFP, respectively. Airyscan images were acquired on a Zeiss LSM980 Observer in SuperResolution mode using a Plan-Apochromat 40x/1.4 Oil objective with excitation lasers of 405 nm, 488 nm and 561 nm wavelengths. Lysotracker signal intensity after LLOMe treatment was measured using FIJI. Gal3, LC3 and TBC1D15 puncta and lysosomal tubules in cells were counted by thresholding the raw images, then converting them to 8-bit images for processing using the “analyse particles” FIJI plugin.For DQ-BSA imaging, images were converted to 8-bit, mean fluorescent intensity in the green channel was measured in the full cell which were marked by LAMP1-RFP.

### Analysis of lysosomal tubules

For lysosomal tubules, particles with a circularity of 0.8–1.0 were filtered out. The length of the remaining tubular structures was determined, and those exceeding 2.5 µm were classified as long tubules. At least 75 cells from three independent experiments were used for statistical analysis. Graphs were plotted in MS Excel. A two-tailed type-3 Student’s t-test was used to determine statistical significance. For image representation labelled as ‘trace’ in the figure, the image was split into separate channels. LAMP1, LC3 and TBC1D15 positive structures were manually traced using the Wand (tracing) tool of FIJI. For STED images, at least 30 cells taken from 3 experiments were analysed in FIJI. Images were converted to 8 bit, thresholded, and analysed by the ‘Analyze particles’ plugin.All lysosomal structures of particle size less than 2µm2 were filtered out. Particles of the size range <2µm2, circularity <0.5 were taken as tubules. Those with higher circularity were classified as rings. The ‘include holes’ option in the ‘Analyze particles’ was used to distinguish between open rings and closed ones.

### STED microscopy

Stimulated Emission Depletion Microscopy (STED) images were taken with an *abberior Instruments* STEDYCON STED setup equipped with an inverted IX83 microscope (Olympus), a 100x oil objective (UPLXAPO100XO, 100x / NA 1.45, oil, Olympus), using pulsed excitation lasers at 640 nm (to excite the *abberior* STAR RED labelled LAMP1), a pulsed STED laser operating at 775 nm, continuous autofocus and gated detection with avalanche photodiode element detectors (APDs). All acquisition operations were controlled by the STEDYCON Software (abberior Instruments).

### Proximity labelling experiments

All samples were processed as three biological replicates and proximity labelling experiments were based on the doxycycline-inducible expression of APEX2-tagged gene constructs. Cells expressing APEX2-LAMP1, APEX2-TBC1D15 or APEX2-TRPML1 were induced with 0.5μg/ml doxycycline for 24 h before each treatment, and we added 500μM of biotin-tyramide for the final hour. We added 1 mM H_2_O_2_ for 1 min to trigger the biotinylation of nearby substrates. The cells were then washed three times with PBS containing10 mM sodium azide, 10 mM sodium ascorbate and 5 mM Trolox to quench the reaction, followed by a PBS wash to remove the chemicals. The cells were lysed (20 mM HEPES-KOH pH 7.5, 150 mM KCl, 0.2 mM EDTA, 0.5% NP-40) and 1 mg of the lysate was incubated with 20μl streptavidin-agarose resin overnight at 4 °C. The immunoprecipitated biotinylated proteins were washed five times in lysis buffer followed by western blotting as described above, or another five washes in MS-grade water prior to MS analysis.

### Mass spectrometry

Washed immunoprecipitated samples were mixed with 40 M urea for 3 h at 37 °C, then ttthe proteins were reduced with 1 mM TCEP and alkylated with 4 mM chloroacetamide for 1 h. The samples were then diluted in 50 mM ammonium bicarbonate to reduce the urea concentration below 1 M and digested with 0.5μg Lys-c and 1μg trypsin at 37 °C for 14–18 h. The digested samples were acidified with 1% trifluoroacetic acid and the peptides were desalted using either Sep-Pak cartridges or C-18 stage tips. Dried peptides were resuspended in TMT buffer and labelled with 6-plex TMT for MS analysis as previously described (Shin et al., 2021). All samples were processed as at least three biological replicates.

Samples were analysed on a Q Exactive HF coupled to an easy nLC 1200 (ThermoFisher Scientific) using a 35 cm long, 75µm ID fused-silica column packed in house with 1.9 µm C18 particles (Reprosil pur, Dr. Maisch), and kept at 50°C using an integrated column oven (Sonation). TMT6 labelled peptides were eluted by a non-linear gradient from 4-32% acetonitrile over 120 minutes and directly sprayed into the mass-spectrometer equipped with a nanoFlex ion source (ThermoFisher Scientific). Full scan MS spectra (350-1400 m/z) were acquired in Profile mode at a resolution of 120,000 at m/z 200, a maximum injection time of 100 ms and an AGC target value of 3 x 10^6^ charges. Up to 20 most intense peptides per full scan were isolated using a 1.0 Th window and fragmented using higher energy collisional dissociation (normalised collision energy of 35). MS/MS spectra were acquired in centroid mode with a resolution of 15,000, a maximum injection time of 50 ms and an AGC target value of 1 x 10^5^. Single charged ions, ions with a charge state above 2 and ions with unassigned charge states were not considered for fragmentation and dynamic exclusion was set to 20s.

IP peptides from label free experiments were eluted by a non-linear gradient from 4-32% acetonitrile over 60 minutes and directly sprayed into the mass-spectrometer equipped with a nanoFlex ion source (ThermoFisher Scientific). Full scan MS spectra (300-1650 m/z) were acquired in Profile mode at a resolution of 60,000 at m/z 200, a maximum injection time of 20 ms and an AGC target value of 3 x 10^6^ charges. Up to 10 most intense peptides per full scan were isolated using a 1.4 Th window and fragmented using higher energy collisional dissociation (normalised collision energy of 27). MS/MS spectra were acquired in centroid mode with a resolution of 30,000, a maximum injection time of 54 ms and an AGC target value of 1 x 10^5^. Single charged ions, ions with a charge state above 2 and ions with unassigned charge states were not considered for fragmentation and dynamic exclusion was set to 20s.

Raw MS data were analysed with Proteome Discoverer v2.4 (Thermo Fisher Scientific) using Sequest HT as the search engine and performing re-calibration of precursor masses with the Spectrum RC-node. Fragment spectra were screened against the human reference proteome and against common contaminants in ‘contaminants.fasta’ provided with MaxQuant. The accepted static modifications were TMTs on the N-terminus and lysine side chains as well as carbamidomethylated cysteine residues. Accepted dynamic modifications were methionine oxidation and N-terminal acetylation. Matched spectra were filtered with Percolator, applying an FDR of 1% on the peptide spectrum match and protein level. Reporter intensities were normalized to the total protein intensities in Proteome Discoverer, assuming equal sample loading, and also by median normalization using the NormalyzerDE package, if required (Willfross et al., 2018). Label-free data were analysed using MaxQuant v1.65 (Cox and Mann, 2008). Fragment spectra were screened against the *Homo sapiens* SWISSPROT database (TaxID: 9606). Label-free quantification was achieved using MaxLFQ (Cox et al., 2014) with activated matches between runs. Statistically significant changes between samples were determined in Perseus v1.6.6.0 based on a threshold of *p* ≤0.01 and a log _2_ fold change threshold of ±0.5 (Tyanova et al., 2016).

Raw MS data were analysed with Proteome Discoverer v2.4 (Thermo Fisher Scientific) using Sequest HT as the search engine and performing re-calibration of precursor masses with the Spectrum RC-node. Fragment spectra were screened against the human reference proteome and against common contaminants in ‘contaminants.fasta’ provided with MaxQuant.The accepted static modifications were TMTs on the N-terminus and lysine side chains as well as carbamidomethylated cysteine residues. Accepted dynamic modifications were methionine oxidation and N-terminal acetylation. Matched spectra were filtered with Percolator, applying an FDR of 1% on the peptide spectrum match and protein level. Reporter intensities were normalized to the total protein intensities in Proteome Discoverer, assuming equal sample loading, and also by median normalization using the NormalyzerDE package, if required (Willfross et al., 2018). Label-free data were analysed using MaxQuant v1.65 (Cox and Mann, 2008). Fragment spectra were screened against the *Homo sapiens* SWISSPROT database (TaxID: 9606). Label-free quantification was achieved using MaxLFQ (Cox et al., 2014) with activated matches between runs. Statistically significant changes between samples were determined in Perseus v1.6.6.0 based on a threshold of *p* ≤ 0.01 and a log_2_ fold change threshold of ±0.5 (Tyanova et al., 2016).

## Acknowledgements

We thank Abberior Instruments GmbH, specifically Dirk Luchtman for performing STED imaging, Oliver Florey (Babraham Institute, UK) for his gift of ATG16L plasmids (WT and K490A). We thank the quantitative proteomics Unit of IBC2 for the use of their proteomics platform and the Frankfurt Center for Advanced Microscopy (FCAM) for access to microscopes. acknowledges funding from the Deutsche Forschungsgemeinschaft (DFG; German Research Foundation) project number 259130777–SFB 1177, the European Research Council (ERC) under the European Union’s Horizon 2020 research and innovation program (grant agreement number 742720), Else Kröner Fresenius Stiftung, Dr. Rolf M. Schwiete Stiftung, and the Ernst Jung Prize for Medicine. A.B. acknowledges funding from LYSOFOR2625. R.M. received funding through an Alexander von Humboldt Stiftung postdoctoral fellowship.

## Author contributions

**AB** and **RM** conceptualized and performed all cell biology and proteomics experiments. **SK** performed structural and sequence analysis, **RR** handled the MS machine runs, **MB and CM** established the MS protocol for the analysis of proximity labelling assays, **RM** and **SJ** performed airy-scan confocal imaging, **AB, RM** and **ID** analysed the data **ID** supervised the project. **AB, RM** and **ID** wrote the manuscript.

## Supplementary figure legends

**Supplementary Figure 1:**
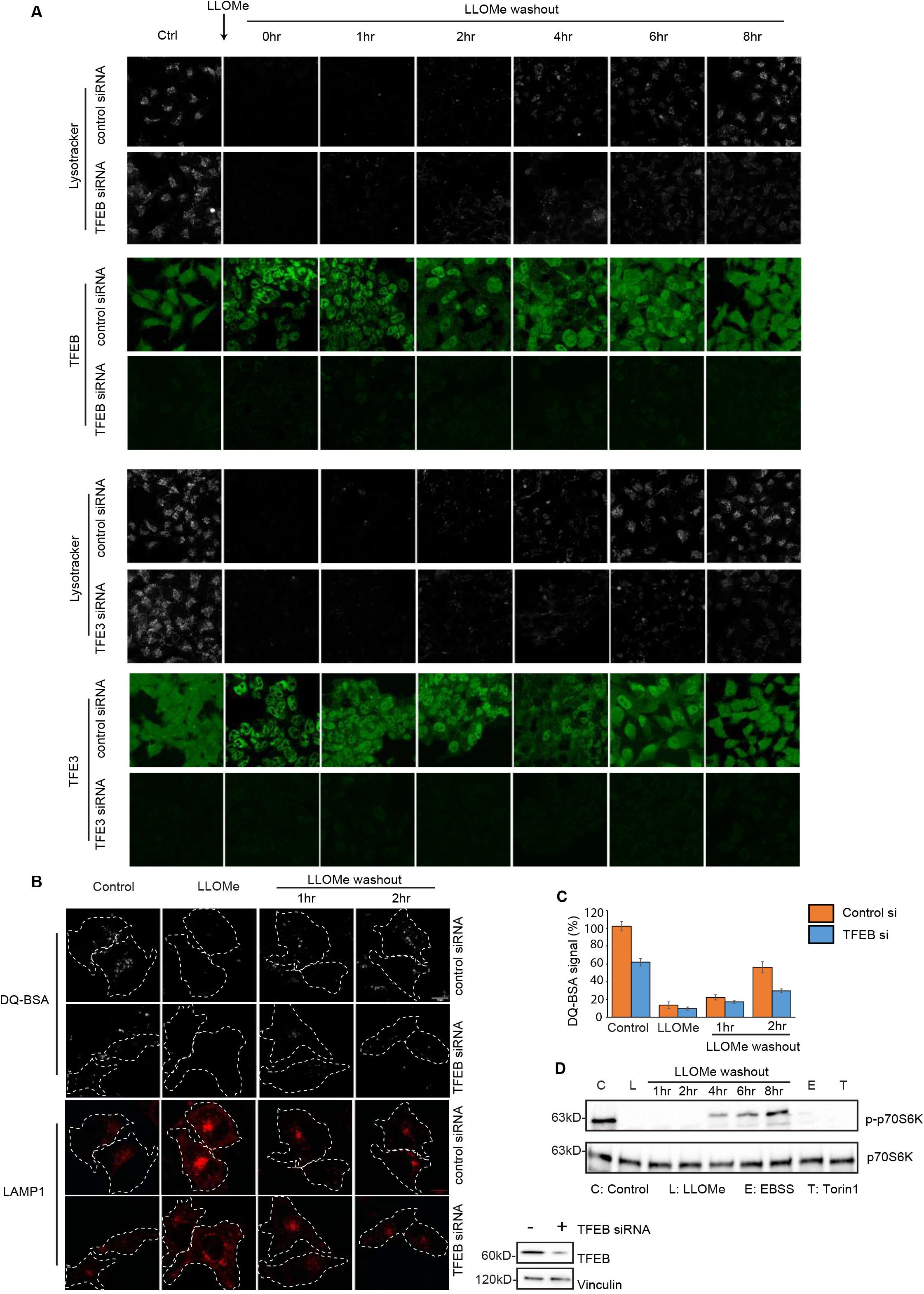
Role of TFEB, TFE3, and mTOR in lysosomal regeneration following LLOMe-mediated damage. a) HeLa cells were treated with siRNA to deplete endogenous TFEB/TFE3. Lysosomal damage was induced with 1mM LLOMe treatment for 2 h followed by washout for the indicated durations. Cells were loaded with LysoTracker Red for 30 min, fixed, and stained with a TFEB/TFE3 specific antibody. b) HeLa cells were treated with siRNA to deplete endogenous TFEB and LAMP1-RFP was transfected to mark lysosomes. These cells were then subjected to LLOMe treatment (1mM, 2 h) followed by washout for indicated timepoints. DQ-BSA was loaded for the last 30 minutes of different treatment conditions prior to imaging the samples by confocal microscopy. TFEB knockdown efficiency was checked by western blotting c) DQ-BSA signal intensities were compared between control cells and cells depleted for endogenous TFEB as a percentage of the signal observed in control siRNA treated cells without LLOMe. Data are means ± SEM from at least 30 cells taken from three independent experiments. d) mTOR activity was assessed by monitoring the phosphorylation status of p70-S6K by western blotting for indicated treatments. LLOMe:1mM, 2 h, EBSS: 4 h, Torin1: 500nM, 6 h.

**Supplementary Figure 2:**
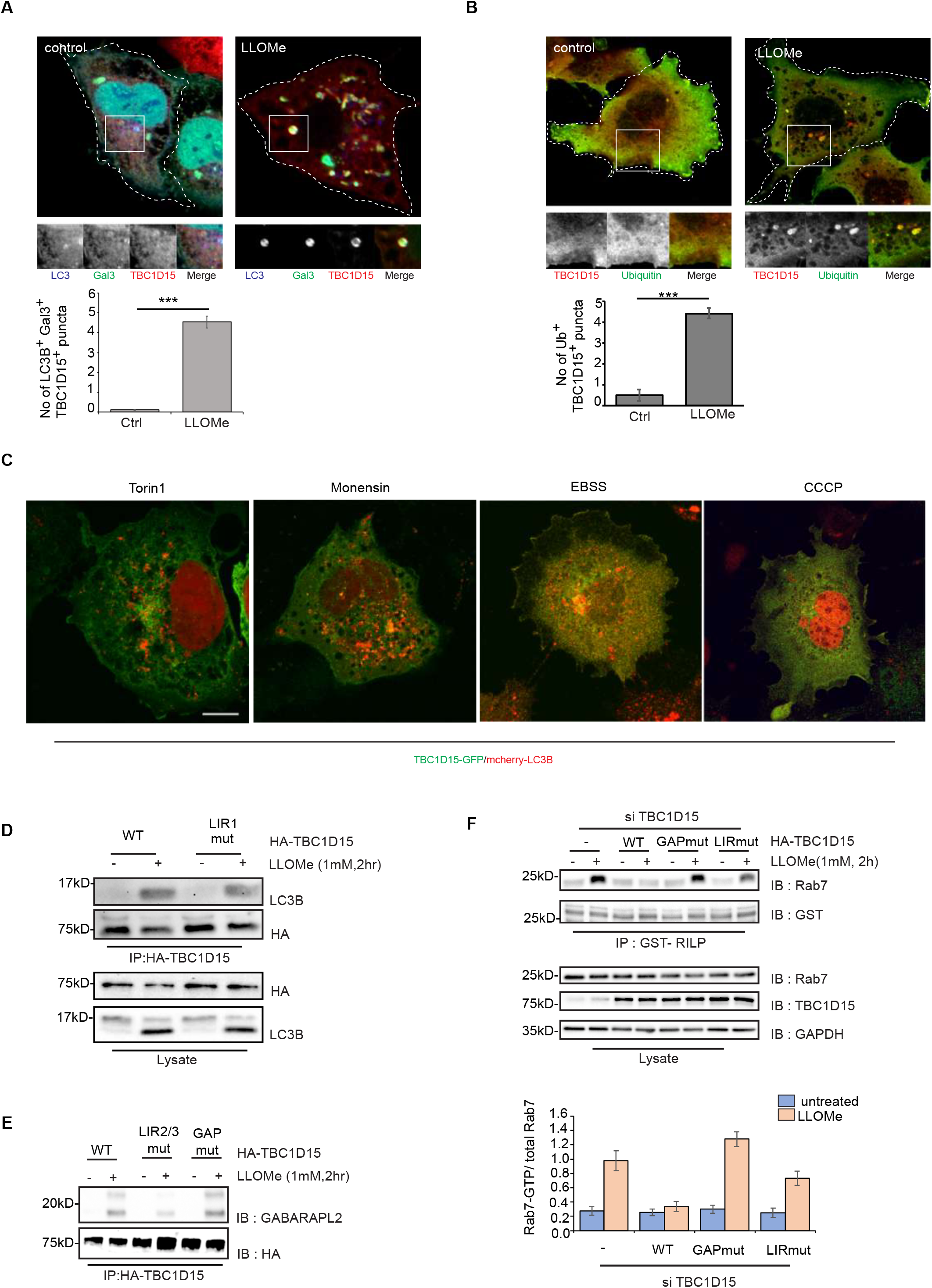
TBC1D15 is recruited to damaged lysosomes following LLOMe treatment but does not colocalize with LC3 under other forms of stress. a) HeLa cells expressing LC3-BFP and GAL3-GFP were treated with 1mM LLOMe for 2 h to induce lysosomal damage. Endogenous TBC1D15 was stained to evaluate its recruitment to the damaged lysosomes. Data are means ± SEM of 50 cells (***p≤ 0.001). b) HeLa cells expressing HA-ubiquitin were treated with LLOMe and endogenous TBC1D15 was stained with an HA-specific antibody to determine the the number of Ub^+^ TBC1D15^+^ puncta/cell. Data are means ± SEM of 50 cells (***p≤0.001). c) HeLa cells expressing TBC1D15-GFP and mCherry-LC3B were treated with Torin1 (500nM, 6 h), Monensin (100µM, 1 h), EBSS (4 h) and CCCP (10µM, 4 h) to induce macroautophagy, non-canonical autophagy (inducer of CASM), macroautophagy and mitophagy respectively. TBC1D15 did not form puncta under these conditions. CASM: <u>C</u>onjugation of <u>A</u>TG8 to <u>s</u>ingle <u>m</u>embranes. d) HEK 293T cells were transfected with wild-type or LIR1 mut variant of HA tagged TBC1D15. Following 1mM LLOMe treatment for 2 h, HA immunoprecipitation was performed to assess the binding with LC3B by western blotting. e) HEK 293T cells were transfected with different HA tagged variants of TBC1D15, as indicated in the figure. HA immunoprecipitation was performed after LLOMe treatment to assess the binding with GABARAPL2 by western blotting. The lysates used here are the same as those shown in Figure 2g. f) HeLa cells were treated with siRNA for 48 h to deplete endogenous TBC1D15 followed by reconstitution with HA-tagged wild-type or mutant TBC1D15 for 24 h. The cells were then treated with LLOMe for 2 h followed by lysis. GTP-bound Rab7 was immunoprecipitated from lysates by incubating with GST-RILP. Total Rab7 levels were assessed by western blotting cell lysates with a Rab7-specific antibody.

**Supplementary Figure 3:**
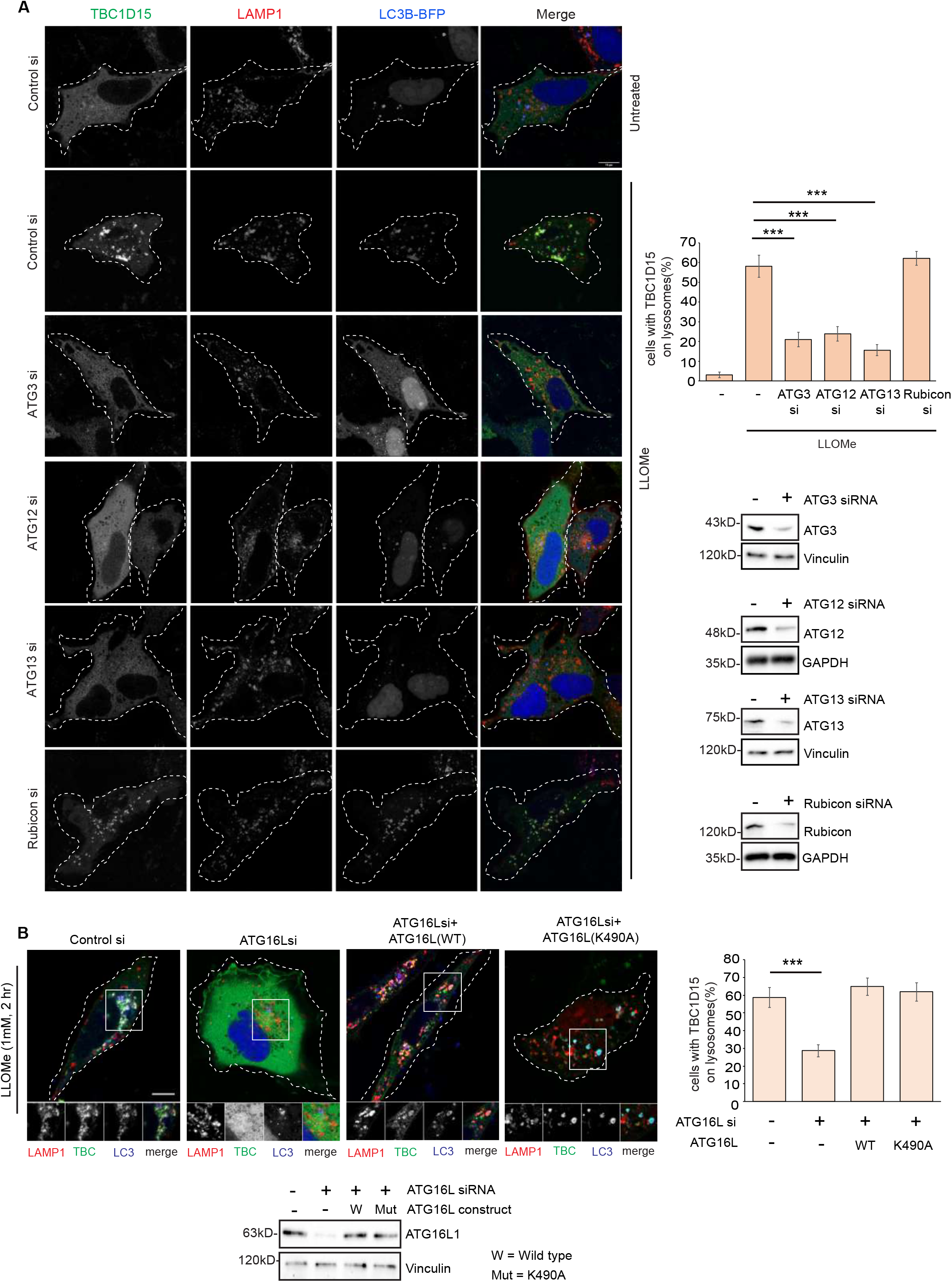
Macroautophagy is important for the recruitment of TBC1D15 to damaged lysosomes. a) HeLa cells were treated with specific siRNA for 48 h to knockdown different autophagy related genes followed by transfection of LC3B-BFP for another 18 hours. Cells were then treated with 1mM LLOMe for 2 h, fixed and stained for LAMP1 and TBC1D15 followed by confocal imaging. Images were analysed in FIJI to determine the number of cells where TBC1D15 is localised on lysosomes. Data are means ± SEM of at least 100 cells taken from three independent experiments (***p≤0.001). Knockdown efficiency for each gene was checked by western blotting. b) HeLa cells were treated with siRNA to knockdown ATG16L followed by reconstituting with either wild-type or K490A mutant variants. Cells were then transfected with LC3B-BFP for 18 h, followed by 1mM LLOMe treatment for 2 h and stained for LAMP1 and TBC1D15 to quantitate the recruitment of TBC1D15 to damaged lysosomes. Data are means ± SEM of at least 80 cells from three independent experiments (***p≤0.001). Expression levels of ATG16L was checked by western blotting.

**Supplementary Figure 4:**
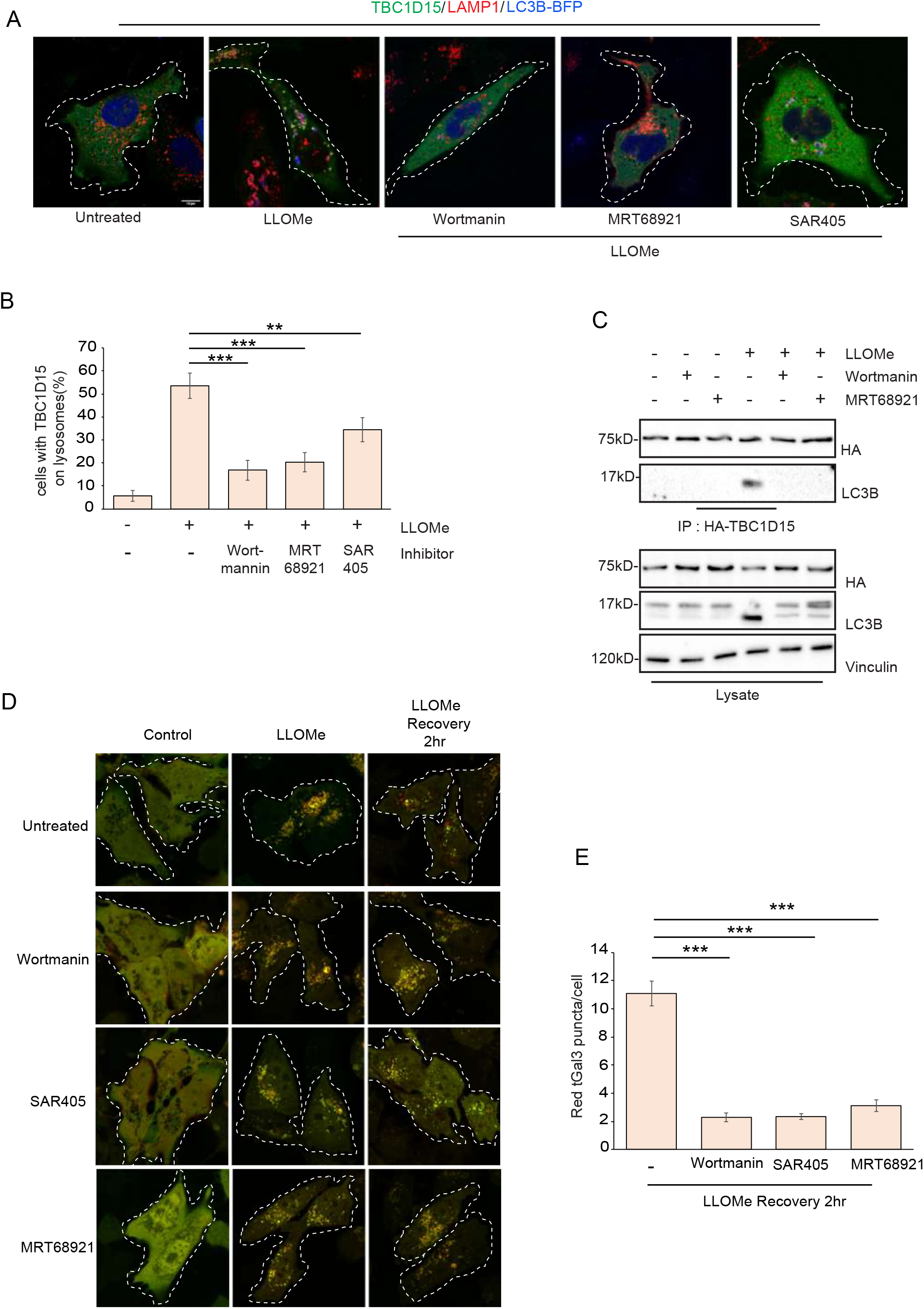
Pharmacological inhibition of autophagy reduces recruitment of TBC1D15 to damaged lysosomes and reduces lysosomal regeneration flux. a) HeLa cells were transfected with HA-TBC1D15 and LC3B-BFP followed by treating them with only 1mM LLOMe for 2 h or LLOMe in combination with different known chemical agents to block autophagy. Cells were then fixed and stained for LAMP1 to check the recruitment of TBC1D15 to damaged lysosomes. b) Recruitment efficiency of TBC1D15 from experiment in panel (a) Data are means ± SEM of at least 80 cells from three independent experiments (***p≤0.001, **0.001 < p≤ 0.01). c) HEK 293T cells were transfected with HA tagged wild-type TBC1D15 followed by treating them with 1mM LLOMe for 2 h or LLOMe in combination with indicated inhibitors.Binding efficiency of TBC1D15 with LC3B under these conditions were assessed by HA immunoprecipitation followed by western blotting. d) Lysosomal regeneration flux assay was performed using HeLa cells stably expressing the tfGal3 reporter after 1mM LLOMe treatment for 2 h followed by LLOMe washout. The indicated drugs were added to the medium 2 h prior to LLOMe treatment, during LLOMe treatment and washout. e) Efficiency of lysosomal regeneration was compared between treatments as shown in experiment in panel (d) by measuring number of red galectin3 puncta/cell. Data are means ± SEM of at least 50 cells from three independent experiments (***p≤0.001). Drug treatments used: Wortmannin (300nM, 6 h), MRT68921 (10µM, 6 h), SAR405 (10µM, 6 h).

**Supplementary Figure 5:**
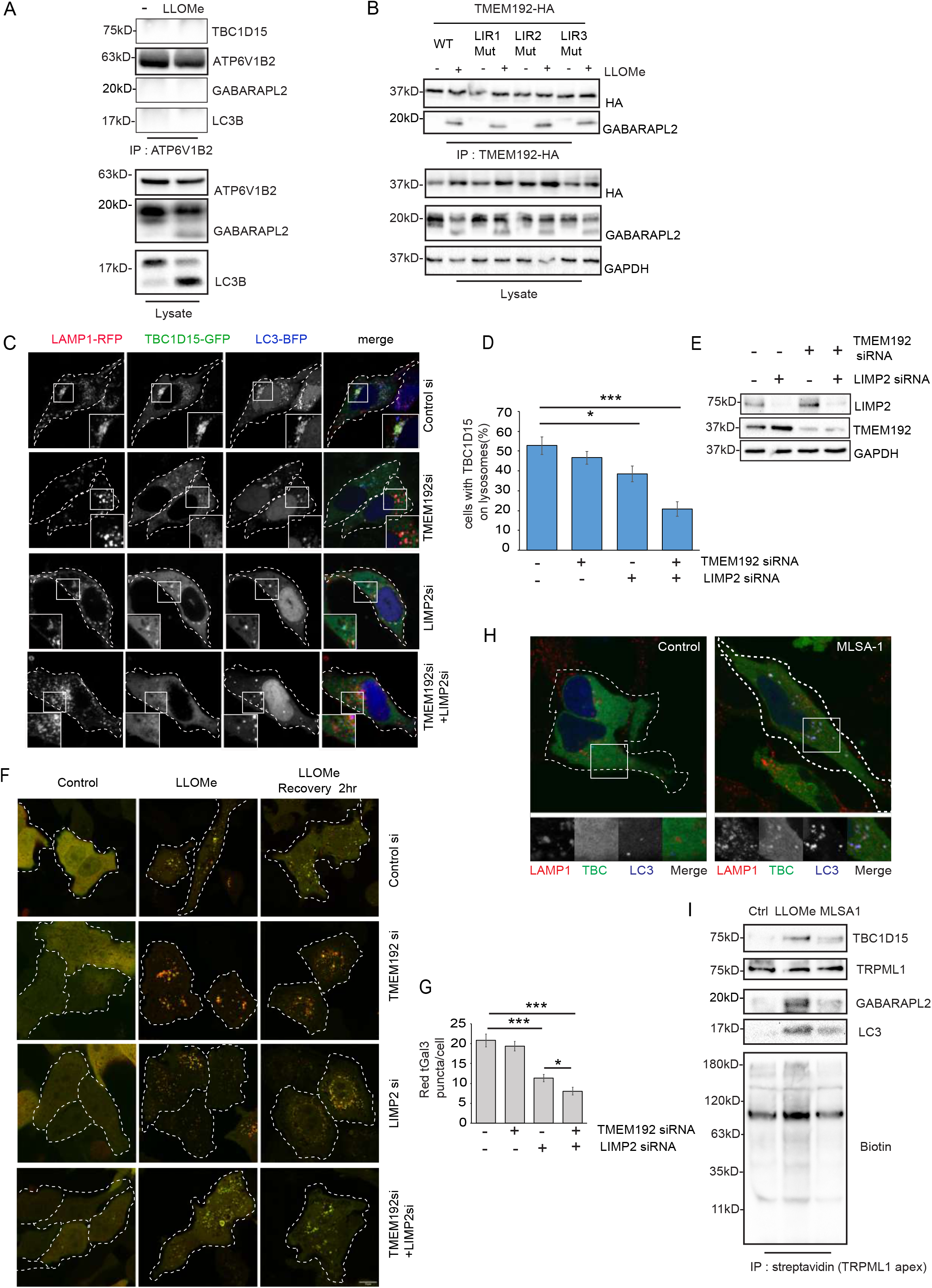
Role of ATG8 binding proteins in TBC1D15 recruitment and lysosomal regeneration. a) Endogenous ATP6V1B2 was immunoprecipitated using an antibody coupled to Protein A/G agarose after treating cells with LLOMe and binding with ATG8 proteins were assessed by western blotting. b) Putative LIR motifs for TMEM192 as described in panel (Fig 4d) were mutated by site directed mutagenesis. HA-tagged TMEM192 constructs were transfected in HEK 293T cells, treated with 1mM LLOMe for 2 h or left untreated. The efficiency of binding with GABARAPL2 for each construct was assessed by HA immunoprecipitation and western blotting. c) HeLa cells were treated with siRNA for 48 h to deplete either LIMP2 and TMEM192 individually or in combination. These cells were then transfected with indicated constructs for 18 h followed by treatment with 1mM LLOMe for 2 h, fixed and imaged by confocal imaging. d) The number of TBC1D15 puncta was compared between control cells and cells treated with siRNA targeting both genes as described in panel (c). Data are means ± SEM of 100 cells from three independent experiments (***p≤0.001; *0.01≤p < 0.05) e) Efficiency of LIMP2 and TMEM192 knockdown were checked by western blotting. f) Importance of LIMP2 and TMEM192 on lysosomal regeneration was assayed by performing the lysosomal regeneration flux assay after LLOMe treatment (1mM, 2 h) and washout (2 h). g) Number of red Gal3 puncta/cell were compared between different samples from experiment in panel (f). Data are means ± SEM of at least 100 cells from three independent experiments (***p≤0.001). h) HeLa cells were transfected with LC3-BFP and were treated with 50 µM MLSA1 for 1 h followed by confocal imaging. TBC1D15 is recruited to LAMP1^+^ LC3^+^ puncta following treatment with MLSA1. i) HeLa cells expressing doxycycline-inducible APEX2-TRPML1 were treated with 1mM LLOMe or 50µM MLSA1 for 2 h followed by proximity labelling using biotin/H_2_O_2_. Lysates were used in a streptavidin pulldown assay followed by western blotting with the indicated antibodies.

**Supplementary Figure 6:**
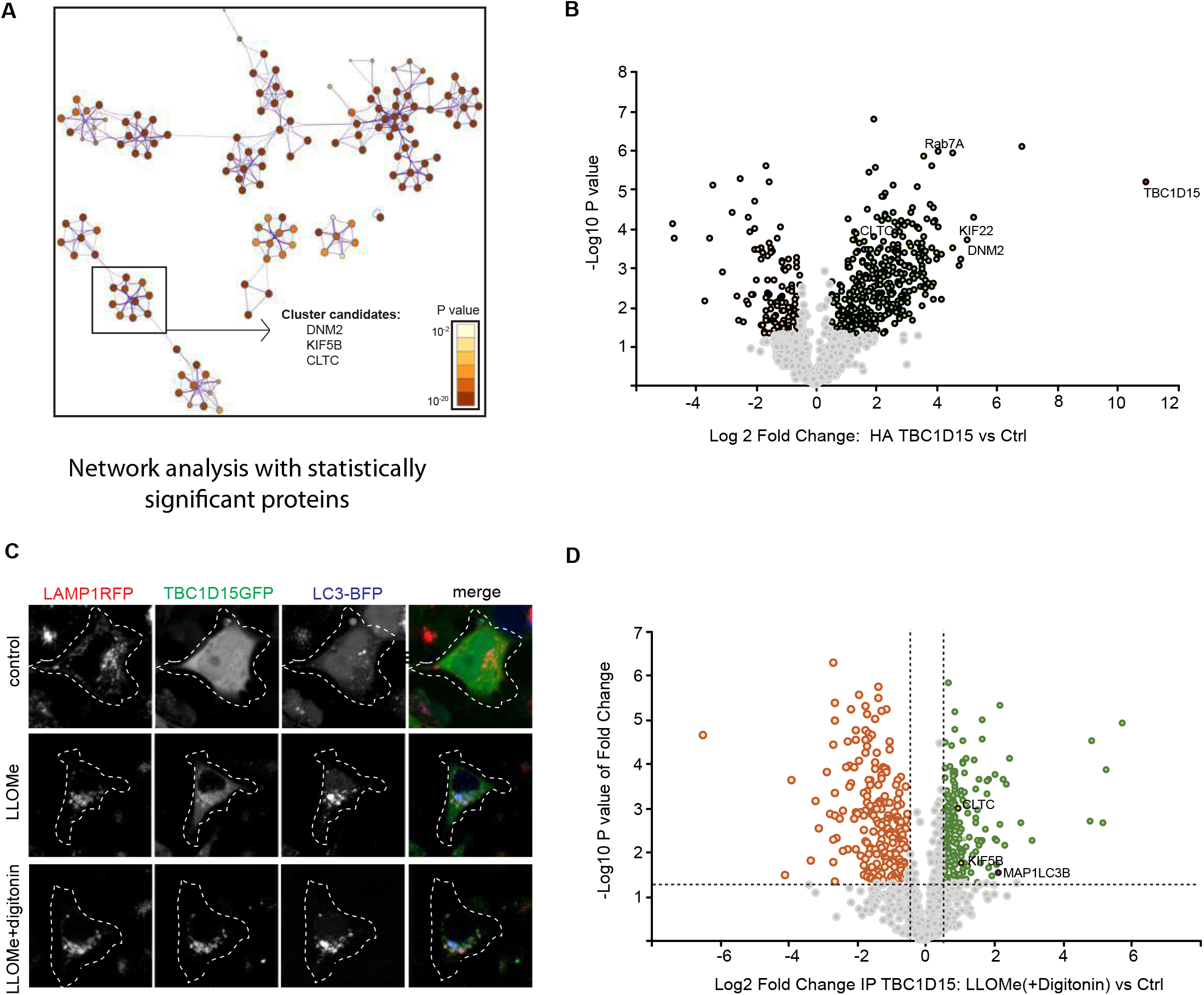
ALR proteins interact with TBC1D15 in LLOMe-treated cells. a) Metascape network analysis of proteins significantly enriched in the proximity labelling experiment shown in Fig 4a and 4b. b) HA-TBC1D15 was immunoprecipitated from cells after 1mM LLOMe treatment for 2 h followed by MS analysis. Proteins related to ALR (KIF22, CLTC and DNM2) were enriched by LLOMe treatment. Control = lysates from cells treated with LLOMe but not expressing HA-TBC1D15. c) Cells expressing LAMP1-RFP, TBC1D15-GFP and LC3-BFP were treated with 1 mM LLOMe for 2 h. Before fixing the cells, 0.05% digitonin was added to reduce the cytosolic background of TBC1D15 staining. Digitonin-treated cells show the presence of TBC1D15 in LAMP1-RFP^+^ LC3-BFP^+^ membrane mass. d) TBC1D15-GFP in lysates from cells treated with LLOMe and digitonin similar to panel (c) was immunoprecipitated using anti-GFP beads followed by MS analysis. ALR proteins were enriched by LLOMe treatment.

**Supplementary Figure 7:**
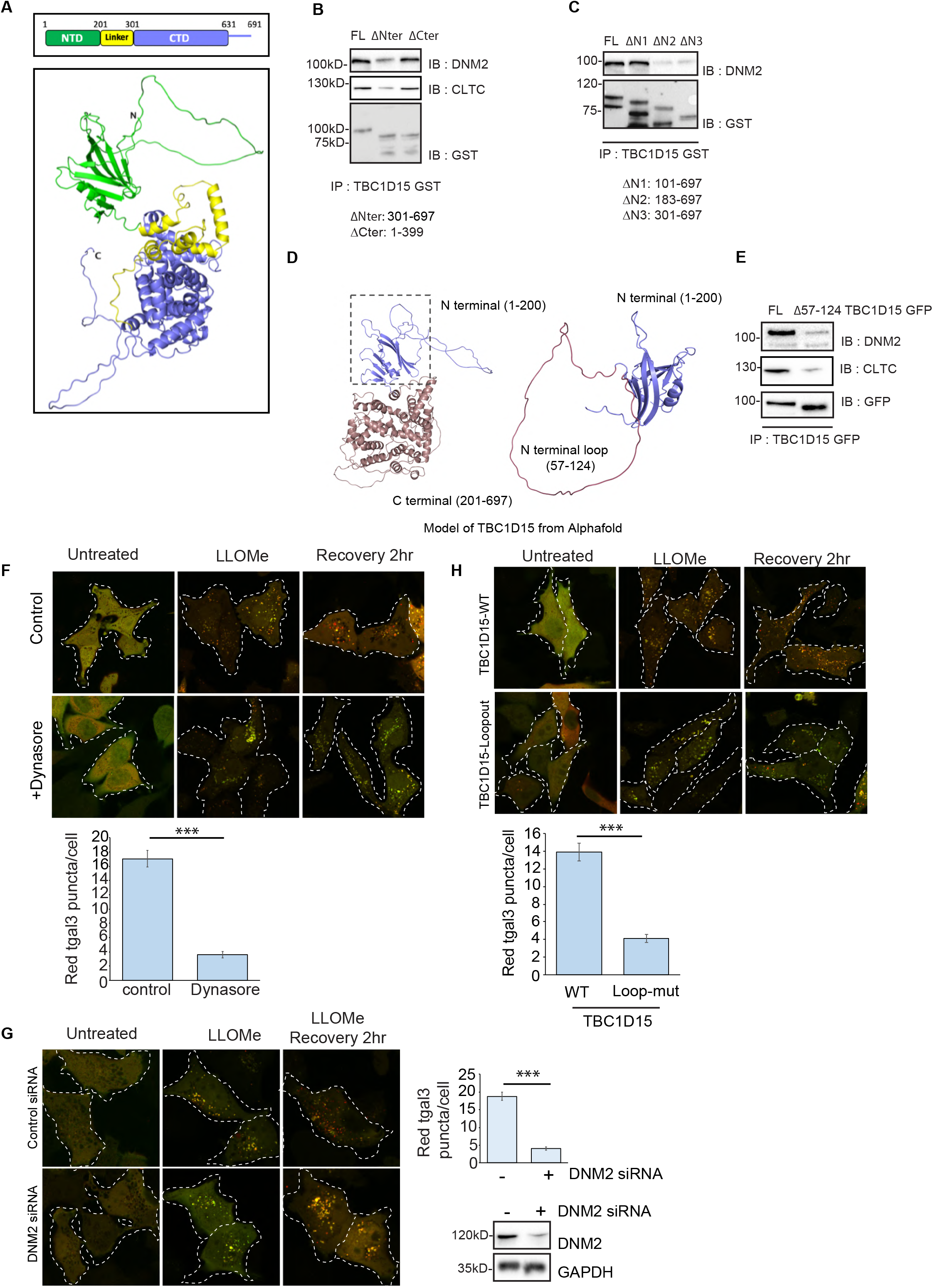
TBC1D15 and DNM2 are important for lysosomal regeneration. a) AlphaFold predicted structure of full-length TBC1D15 showing N-terminal domain (NTD) in green, the linker in yellow and the C-terminal domain (CTD) in blue. b) GST-tagged deletion constructs of TBC1D15 were incubated with cell lysates from HEK 293T cells in a GST pulldown assay and detected by western blotting with the indicated antibodies. c) GST-tagged N-terminal deletion constructs of TBC1D15 were incubated with HEK 293T cell lysates in a GST pulldown assay and detected by western blotting with the indicated antibodies. d) AlphaFold predicted structure showing the N-terminal loop required for DNM2 binding. e) GFP-tagged wild-type TBC1D15 or the loop mutant were enriched from LLOMe-treated cells using anti-GFP beads followed by western blotting with the indicated antibodies. f) HeLa cells stably expressing the tfGal3 flux reporter were subjected to a lysosomal regeneration flux assay in cells with or without 20 µM Dynasore added in LLOMe-free medium during washout. The number of red tGal3 puncta per cell was counted. Data are means ± SEM of at least 50 cells from three independent experiments (***p≤0.001). g) HeLa cells expressing the tfGal3 flux reporter were treated with DNM2 siRNA and then were subjected to a lysosomal regeneration flux assay. The number of red tfGal3 puncta per cell was counted. Data are means ± SEM of at least 50 cells from three independent experiments. (***p≤0.001). h) HeLa cells stably expressing tfGal3 were depleted of endogenous TBC1D15 for 48 h followed by transfection of wild-type TBC1D15 or the loop mutant. Cells were treated with 1mM LLOMe for 2h followed by LLOMe washout and imaged by confocal microscopy. The number of red tGal3 puncta per cell was counted. Data are means ± SEM of at least 50 cells from three independent experiments (***p≤0.001).

**Supplementary Figure 8:**
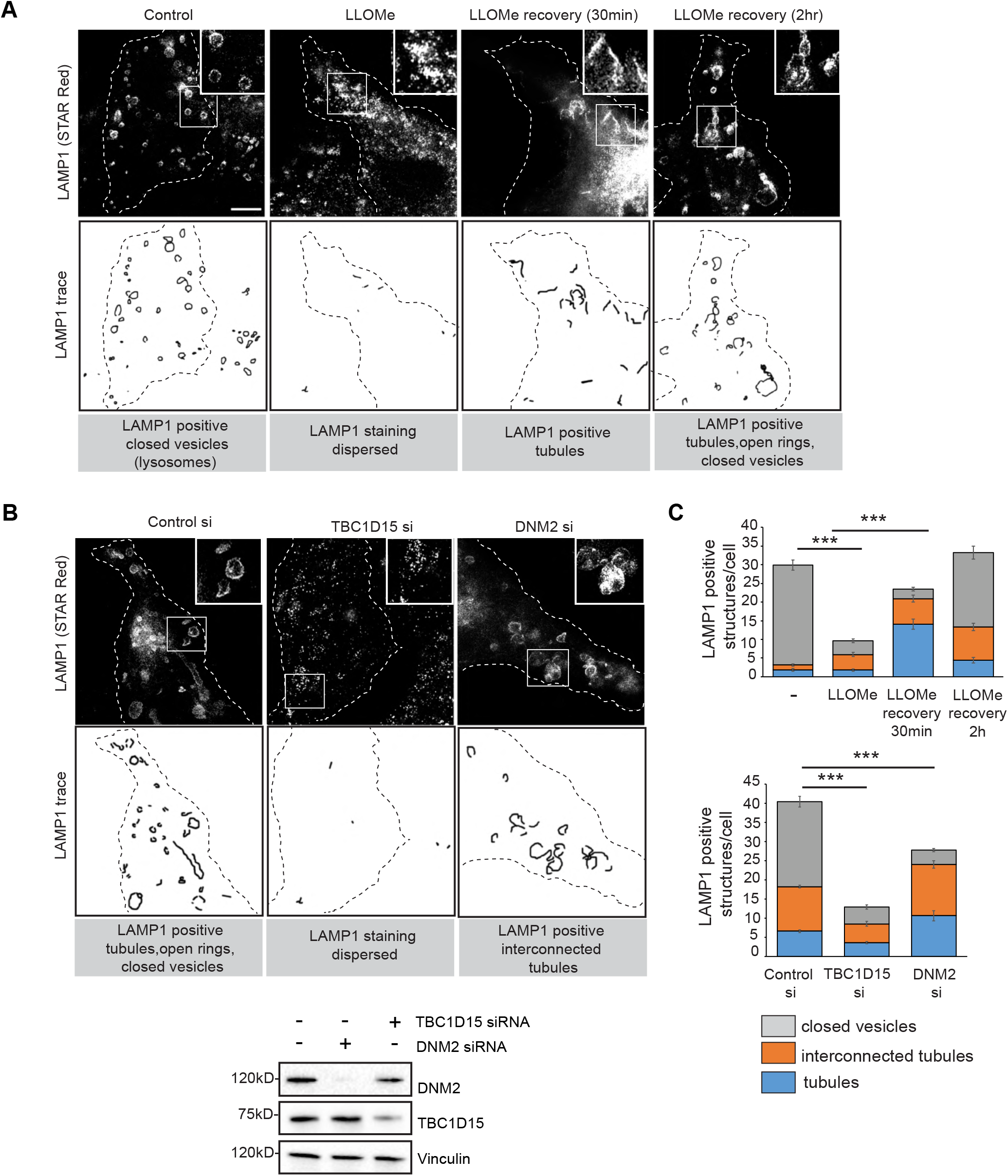
Lysosomal regeneration from damaged membranes occurs via tubulation and successive scission in a TBC1D15 and DNM2 dependent manner. a) HeLa cells were treated with LLOMe and were allowed to recover for the indicated timepoints. Cells were fixed and stained for LAMP1 and imaged by STED microscopy. b) HeLa cells were treated with siRNA to deplete either TBC1D15 or DNM2 individually. These cells were the allowed to recover for 2hr after LLOMe treatment and were then fixed and stained for LAMP1 to assess the effect of the TBC1D15 and DNM2 on lysosomal tubulation events by STED microscopy. c) Different lysosomal structures, from experiments in panel (a) and (b) were analysed in FIJI and classified into closed rings, interconnected tubules and tubules Data are means ± SEM of at least 50 cells from three independent experiments (***p≤0.001).

**Supplementary Figure 9:**
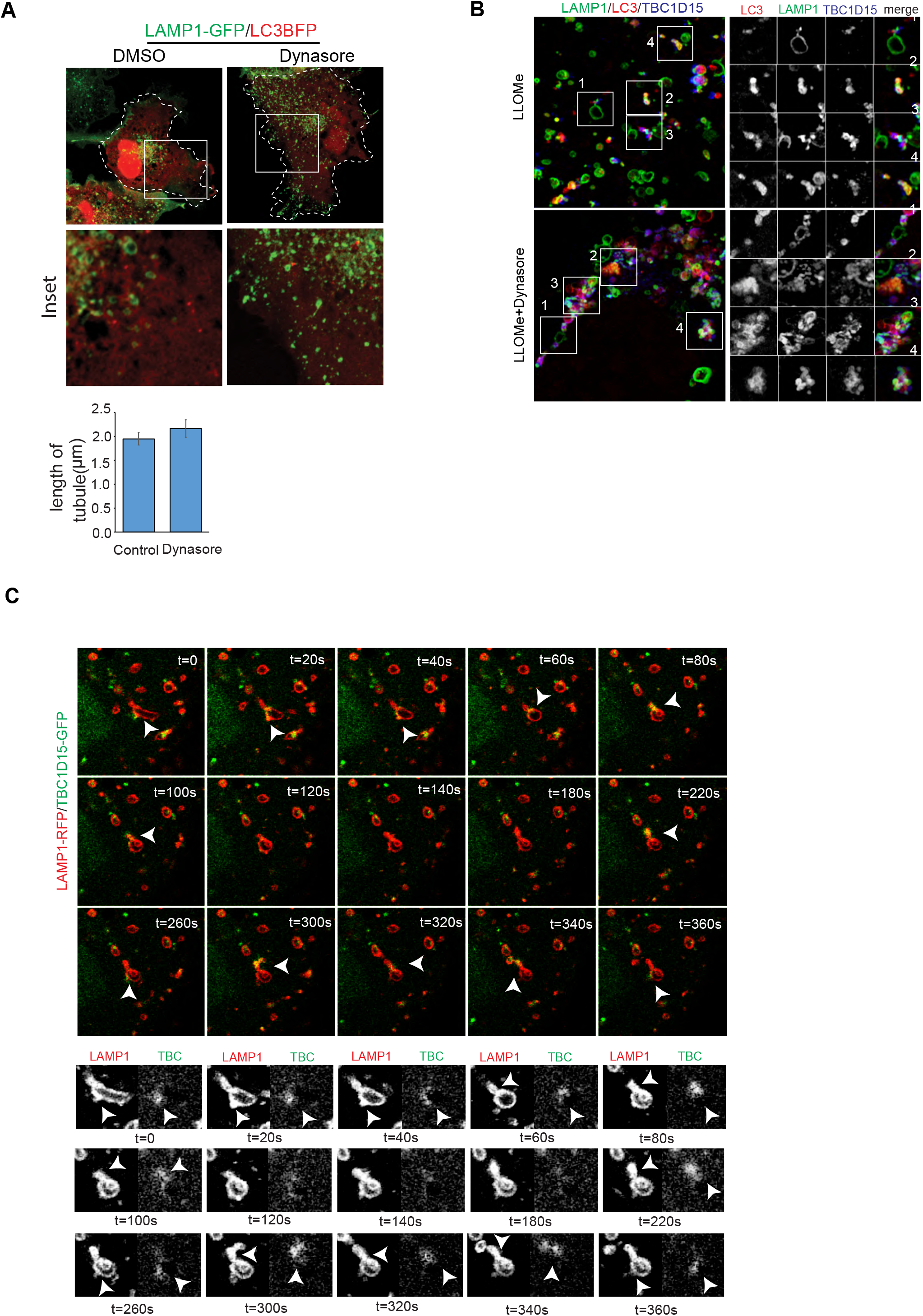
Lysosomal regeneration occurs via tubulation and successive scission in a TBC1D15 and DNM2 dependent manner. a) Effect of 20µM dynasore treatment for 2 h in the absence of LLOMe. b) 3D projections made from Z-stacks of airy-scan images as seen in Fig 5B. c) Time-lapse imaging of 1mM LLOMe treated cells expressing LAMP-RFP and TBC1D15-GFP.

## References

Bonam, S.R., Wang, F. and Muller, S., 2019. Lysosomes as a therapeutic target. Nature Reviews Drug Discovery, 18(12), pp.923–948.

Boya, P. and Kroemer, G., 2008. Lysosomal membrane permeabilization in cell death. Oncogene, 27(50), pp.6434–6451.

Chauhan, S., Kumar, S., Jain, A., Ponpuak, M., Mudd, M.H., Kimura, T., Choi, S.W., Peters, R., Mandell, M., Bruun, J.A. and Johansen, T., 2016. TRIMs and galectins globally cooperate and TRIM16 and galectin-3 co-direct autophagy in endomembrane damage homeostasis. Developmental cell, 39(1), pp.13–27.

Chen, Y. and Yu, L., 2017. Recent progress in autophagic lysosome reformation. Traffic, 18(6), pp.358–361.

Cox, J. and Mann, M., 2008. MaxQuant enables high peptide identification rates, individualized ppb-range mass accuracies and proteome-wide protein quantification. Nature biotechnology, 26(12), pp.1367–1372.

Cox, J., Hein, M.Y., Luber, C.A., Paron, I., Nagaraj, N. and Mann, M., 2014. Accurate proteome-wide label-free quantification by delayed normalization and maximal peptide ratio extraction, termed MaxLFQ. Molecular & cellular proteomics, 13(9), pp.2513–2526.

De Duve, C., Pressman, B.C., Gianetto, R., Wattiaux, R. and Appelmans, F., 1955. Tissue fractionation studies. 6. Intracellular distribution patterns of enzymes in rat-liver tissue. Biochemical Journal, 60(4), p.604.

Dehay, B., Bové, J., Rodríguez-Muela, N., Perier, C., Recasens, A., Boya, P. and Vila, M., 2010. Pathogenic lysosomal depletion in Parkinson’s disease. Journal of Neuroscience, 30(37), pp.12535–12544.

Eapen, V.V., Swarup, S., Hoyer, M.J., Paulo, J.A. and Harper, J.W., 2021. Quantitative proteomics reveals the selectivity of ubiquitin-binding autophagy receptors in the turnover of damaged lysosomes by lysophagy. Elife, 10, p.e72328.

Emmerson, B.T., Cross, M., Osborne, J.M. and Axelsen, R.A., 1990. Reaction of MDCK cells to crystals of monosodium urate monohydrate and uric acid. Kidney international, 37(1), pp.36–43.

Fletcher, K., Ulferts, R., Jacquin, E., Veith, T., Gammoh, N., Arasteh, J. M., … & Florey, O. (2018). The WD 40 domain of ATG 16L1 is required for its non-canonical role in lipidation of LC 3 at single membranes. The EMBO journal, 37(4), e97840.

Fujita, N., Morita, E., Itoh, T., Tanaka, A., Nakaoka, M., Osada, Y., Umemoto, T., Saitoh, T., Nakatogawa, H., Kobayashi, S. and Haraguchi, T., 2013. Recruitment of the autophagic machinery to endosomes during infection is mediated by ubiquitin. Journal of Cell Biology, 203(1), pp.115–128.

Jacomin, A.C., Samavedam, S., Promponas, V. and Nezis, I.P., 2016. iLIR database: A web resource for LIR motif-containing proteins in eukaryotes. Autophagy, 12(10), pp.1945–1953.

Jia, J., Claude-Taupin, A., Gu, Y., Choi, S.W., Peters, R., Bissa, B., Mudd, M.H., Allers, L., Pallikkuth, S., Lidke, K.A. and Salemi, M., 2020. Galectin-3 coordinates a cellular system for lysosomal repair and removal. Developmental cell, 52(1), pp.69–87.

Jumper, J., Evans, R., Pritzel, A., Green, T., Figurnov, M., Ronneberger, O., Tunyasuvunakool, K., Bates, R., Žídek, A., Potapenko, A. and Bridgland, A., 2021. Highly accurate protein structure prediction with AlphaFold. Nature, 596(7873), pp.583–589.

Khundadze, M., Ribaudo, F., Hussain, A., Stahlberg, H., Brocke-Ahmadinejad, N., Franzka, P., Varga, R.E., Zarkovic, M., Pungsrinont, T., Kokal, M. and Ganley, I.G., 2021. Mouse models for hereditary spastic paraplegia uncover a role of PI4K2A in autophagic lysosome reformation. Autophagy, 17(11), pp.3690–3706.

Koerver, L., Papadopoulos, C., Liu, B., Kravic, B., Rota, G., Brecht, L., Veenendaal, T., Polajnar, M., Bluemke, A., Ehrmann, M. and Klumperman, J., 2019. The ubiquitin-conjugating enzyme UBE 2 QL 1 coordinates lysophagy in response to endolysosomal damage. EMBO reports, 20(10), p.e48014.

Maejima, I., Takahashi, A., Omori, H., Kimura, T., Takabatake, Y., Saitoh, T., Yamamoto, A., Hamasaki, M., Noda, T., Isaka, Y. and Yoshimori, T., 2013. Autophagy sequesters damaged lysosomes to control lysosomal biogenesis and kidney injury. The EMBO journal, 32(17), pp.2336–2347.

Martina, J. A., Chen, Y., Gucek, M., & Puertollano, R. (2012). MTORC1 functions as a transcriptional regulator of autophagy by preventing nuclear transport of TFEB. Autophagy, 8(6), 903–914.

Nakamura, S., Shigeyama, S., Minami, S., Shima, T., Akayama, S., Matsuda, T., Esposito, A., Napolitano, G., Kuma, A., Namba-Hamano, T. and Nakamura, J., 2020. LC3 lipidation is essential for TFEB activation during the lysosomal damage response to kidney injury. Nature cell biology, 22(10), pp.1252–1263.

Napolitano, G. and Ballabio, A., 2016. TFEB at a glance. Journal of cell science, 129(13), pp.2475–2481.

Onoue, K., Jofuku, A., Ban-Ishihara, R., Ishihara, T., Maeda, M., Koshiba, T., Itoh, T., Fukuda, M., Otera, H., Oka, T. and Takano, H., 2013. Fis1 acts as a mitochondrial recruitment factor for TBC1D15 that is involved in regulation of mitochondrial morphology. Journal of cell science, 126(1), pp.176–185.

Papadopoulos, C., Kirchner, P., Bug, M., Grum, D., Koerver, L., Schulze, N., Poehler, R., Dressler, A., Fengler, S., Arhzaouy, K. and Lux, V., 2017. VCP/p97 cooperates with YOD 1, UBXD 1 and PLAA to drive clearance of ruptured lysosomes by autophagy. The EMBO journal, 36(2), pp.135–150.

Peralta, E.R., Martin, B.C. and Edinger, A.L., 2010. Differential effects of TBC1D15 and mammalian Vps39 on Rab7 activation state, lysosomal morphology, and growth factor dependence. Journal of Biological Chemistry, 285(22), pp.16814–16821.

Popovic, D., Akutsu, M., Novak, I., Harper, J.W., Behrends, C. and Dikic, I., 2012. Rab GTPase-activating proteins in autophagy: regulation of endocytic and autophagy pathways by direct binding to human ATG8 modifiers. Molecular and cellular biology, 32(9), pp.1733–1744.

Radulovic, M., Schink, K.O., Wenzel, E.M., Nähse, V., Bongiovanni, A., Lafont, F. and Stenmark, H., 2018. ESCRT-mediated lysosome repair precedes lysophagy and promotes cell survival. The EMBO journal, 37(21), p.e99753.

Repnik, U., Borg Distefano, M., Speth, M. T., Ng, M. Y. W., Progida, C., Hoflack, B., … & Griffiths, G. (2017). L-leucyl-L-leucine methyl ester does not release cysteine cathepsins to the cytosol but inactivates them in transiently permeabilized lysosomes. Journal of cell science, 130(18), 3124–3140.

Saftig, P. and Puertollano, R., 2021. How lysosomes sense, integrate, and cope with stress. Trends in biochemical sciences, 46(2), pp.97–112.

Skowyra, M.L., Schlesinger, P.H., Naismith, T.V. and Hanson, P.I., 2018. Triggered recruitment of ESCRT machinery promotes endolysosomal repair. Science, 360(6384), p.eaar5078.

Sun, J., Deghmane, A.E., Bucci, C. and Hmama, Z., 2009. Detection of activated Rab7 GTPase with an immobilized RILP probe. In Macrophages and Dendritic Cells (pp. 57-69). Humana Press.

Thiele, D.L. and Lipsky, P.E., 1990. Mechanism of L-leucyl-L-leucine methyl ester-mediated killing of cytotoxic lymphocytes: dependence on a lysosomal thiol protease, dipeptidyl peptidase I, that is enriched in these cells. Proceedings of the National Academy of Sciences, 87(1), pp.83–87.

Tyanova, S., Temu, T., Sinitcyn, P., Carlson, A., Hein, M.Y., Geiger, T., Mann, M. and Cox, J., 2016. The Perseus computational platform for comprehensive analysis of (prote) omics data. Nature methods, 13(9), pp.731–740.

Willforss, J., Chawade, A. and Levander, F., 2018. NormalyzerDE: online tool for improved normalization of omics expression data and high-sensitivity differential expression analysis. Journal of proteome research, 18(2), pp.732–740.

Wong, Y.C., Ysselstein, D. and Krainc, D., 2018. Mitochondria–lysosome contacts regulate mitochondrial fission via RAB7 GTP hydrolysis. Nature, 554(7692), pp.382–386.

Yamano, K., Fogel, A.I., Wang, C., van der Bliek, A.M. and Youle, R.J., 2014. Mitochondrial Rab GAPs govern autophagosome biogenesis during mitophagy. Elife, 3, p.e01612.

Yoo, S.M. and Jung, Y.K., 2018. A molecular approach to mitophagy and mitochondrial dynamics. Molecules and cells, 41(1), p.18.

Yoshida, Y., Yasuda, S., Fujita, T., Hamasaki, M., Murakami, A., Kawawaki, J., Iwai, K., Saeki, Y., Yoshimori, T., Matsuda, N. and Tanaka, K., 2017. Ubiquitination of exposed glycoproteins by SCFFBXO27 directs damaged lysosomes for autophagy. Proceedings of the National Academy of Sciences, 114(32), pp.8574–8579.

Yu, L., McPhee, C.K., Zheng, L., Mardones, G.A., Rong, Y., Peng, J., Mi, N., Zhao, Y., Liu, Z., Wan, F. and Hailey, D.W., 2010. Termination of autophagy and reformation of lysosomes regulated by mTOR. Nature, 465(7300), pp.942–946.

Zhou, Y., Zhou, B., Pache, L., Chang, M., Khodabakhshi, A.H., Tanaseichuk, O., Benner, C. and Chanda, S.K., 2019. Metascape provides a biologist-oriented resource for the analysis of systems-level datasets. Nature communications, 10(1), pp.1–10.

